# Neuronal recordings in head-fixed and freely-moving mole-rats

**DOI:** 10.64898/2025.12.15.693140

**Authors:** Runita N. Shirdhankar, Alireza Saeedi, Georgina E. Fenton, Leif Moritz, Pierre-Yves Jacob, Leoni M. Webb, Sybille Wolf-Kümmeth, Wolfgang Boenigk, Emilia Heinrichs, Sabine Begall, E. Pascal Malkemper

## Abstract

Mole-rats are subterranean rodents that have evolved remarkable sensory adaptations to life in underground tunnel systems, yet their neural mechanisms remain largely unexplored. Here, we present a protocol for in vivo electrophysiological recordings in awake, head-fixed, and freely moving African mole-rats (*Fukomys anselli/micklemi*), overcoming unique challenges of studying the neurobiology of subterranean species. For example, we find that mole-rat brain physiology impacts survival after surgeries, with higher carbon dioxide concentrations required for recovery compared to other rodents, likely due to a mutation in the chloride-potassium symporter KCC2. Having addressed the challenges, we used tetrodes and Neuropixels probes to record single-unit activity and local field potentials (LFP) across several cortical and subcortical regions for several weeks. We observed single units responsive to auditory and visual stimuli in the superior colliculus, and hippocampal recordings in freely moving mole-rats revealed prominent theta rhythms at frequencies lower than those observed in any other rodent species to date. Finally, we performed integrated three-dimensional and two-dimensional probe-track analysis within the same brain using tissue clearing, light sheet imaging, rehydration, and vibratome sectioning, and we present a newly developed stereotaxic brain atlas for implantation and histological alignment. The established methodology will guide future studies in comparative rodent neurobiology, providing further insights into neurobiological adaptations to subterranean environments. Given their phylogenetic and ecological similarities, we expect our protocols to be transferable to other subterranean species, including the widely studied naked mole-rat (*Heterocephalus glaber*).

**Highlights:**

- Protocols for chronic and acute electrode implantations in mole-rats
- Stereotaxic brain atlas for the Ansell’s mole-rat (https://doi.org/10.17617/3.UNDKRO)
- Neuropixels and tetrode single-unit recordings in head-fixed and freely moving mole-rats
- Integrated 3D (tissue clearing) and 2D (histology) probe-track analysis within the same brain
- Discovery of low-frequency hippocampal theta rhythm in Ansell’s mole-rats

## Introduction

Since the 1970s, neuroscience has predominantly focused on model species like mice, flies, and zebrafish to unravel neural mechanisms and develop cutting-edge methodologies. While this approach has provided invaluable insights, its narrow scope limits generalizability. Furthermore, some questions in neuroscience such as the functional organization of special sensory modalities, e.g. electric, magnetic, or infrared senses, cannot be answered with the established animal models. Consequently, there is a growing need to explore a broader range of organisms and this trend is enabled by latest technological developments (Brenowitz & Zakon, 2015; Striedter et al., 2014; Yartsev, 2017). Studying diverse species can illuminate unique adaptations and broaden our understanding of the evolutionary landscape of neural processes. African mole-rats (family Bathyergidae) exemplify this opportunity. The last common ancestor of bathyergids and murids like mice and rats, the most-established rodent models in neuroscience, dates back to the Cretaceous more than 70 mya (Figure 1). Over this long time, (social) African mole-rats have acquired extraordinary adaptations to their underground habitats, including tolerance to extreme hypoxia (Park et al., 2017), complex social structures (Kverková et al., 2018), and specialized sensory systems (Vice et al., 2021). Their reduced visual (Němec et al., 2008) and auditory (Pyott et al., 2020) systems and concurring reliance on other sensory modalities such as a highly developed sense of touch (Catania & Remple, 2002) and the ability to sense the Earth’s magnetic field (Burda et al., 1990), make them interesting study subjects that stand out from other rodents. Among the over 30 species, the naked mole-rat (*Heterocephalus glaber)* has already emerged as a model system for biomedical studies (Buffenstein et al., 2021), and the broader family has great potential for studying the neural basis of unique behavioural and physiological traits.

**Figure 1:**
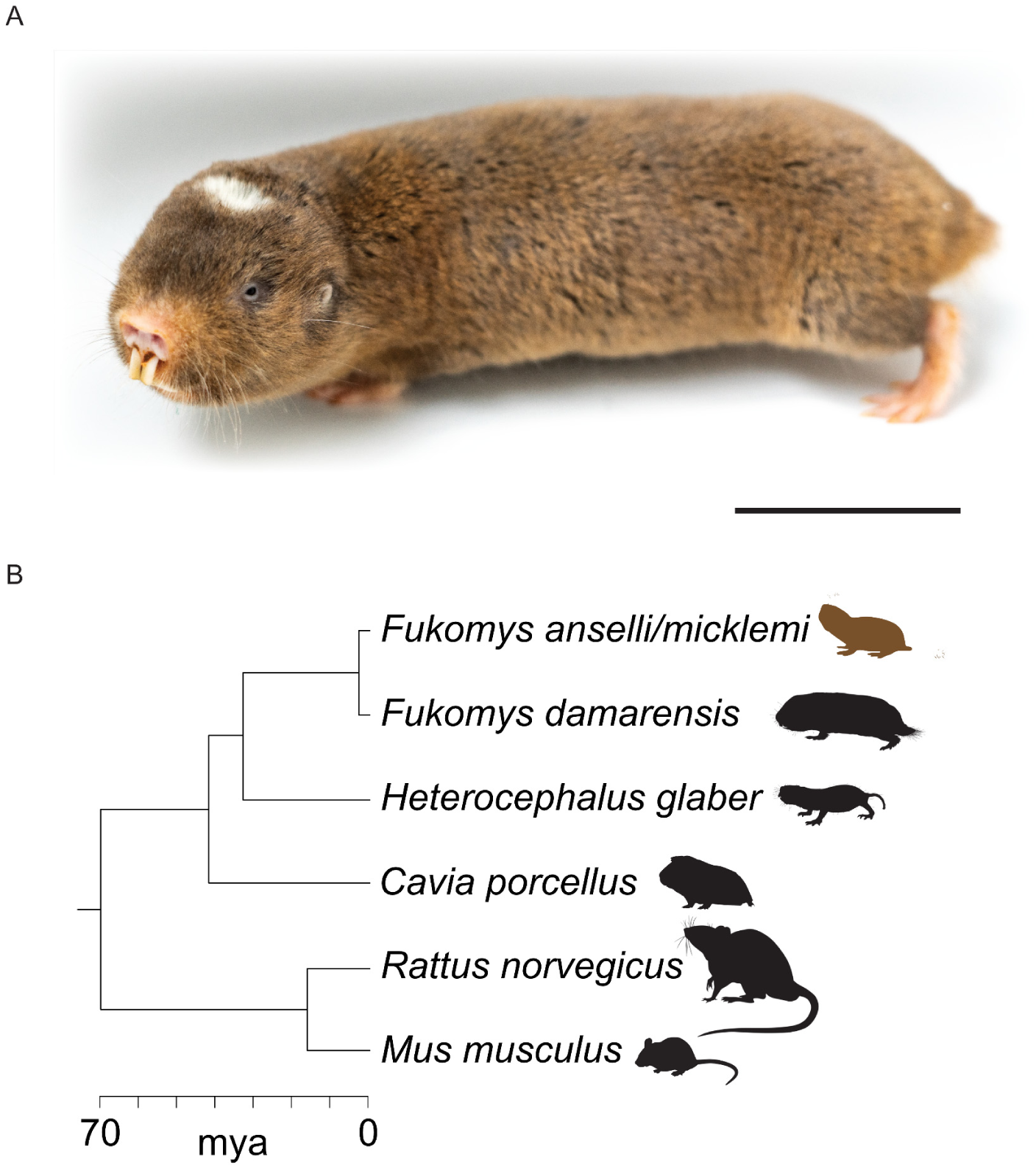
Phylogenetic relationship and divergence time estimates for the studied African mole-rat species and common rodent model organisms. **A.** The model species *Fukomys anselli/micklemi.* The scale bar represents approximately 3 cm. **B.** Phylogenetic tree highlighting the early split between the ancestor of the Hystricomorpha (including African mole-rats and the guinea pig) and the most common rodent model-organisms in neuroscience, rats and mice, 70 million years ago (mya). The guinea pig (*Cavia porcellus*) belongs to the Caviomorpha (New World Hystricognathi), the sister group of African Phiomorpha (Old World Hystricognathi), and can be considered as a neuroscience model organism. The phylogeny and divergence time estimates are based on Afzali & López-Antonanzas (2024) and Voloch et al. (2013) for *Mus* and *Rattus*, and on Pyott et al. (2020) for remaining species. *Fukomys anselli* and *Fukomys micklemi* are shown as a single species here, in line with most recent phylogenetic analyses (Šumbera, pers. comm.). Silhouette images were obtained from PhyloPic (http://phylopic.org) under a public domain license (CC0 1.0 license), except for *Fukomys anselli/micklemi* (<u>credit</u> to Kai Caspar, modified), *Fukomys damarensis* (<u>credit</u> to Kyle Finn), and *Cavia porcellus* (<u>credit</u> to Ricardo Araújo).

Electrophysiology provides unparalleled temporal resolution for studying neuronal activity without requiring genetic modifications, thus is ideally suited to study the brains of species that are not commonly used as models. However, applying this method to non-model species is often challenged by distinct anatomy, physiology, and behavior. Here, we introduce protocols for electrophysiological recordings in both awake, head-fixed, and freely moving African mole-rats (*Fukomys micklemi/anselli*). We uncover critical components of mole-rat physiology that necessitate protocol adaptation to ensure successful surgery. We created a stereotaxic brain atlas and successfully recorded electrophysiological data from the superior colliculus and hippocampus. Our findings highlight the importance of considering the ecology of a species when establishing protocols for non-model organisms and pave the way to study the neuronal mechanisms underlying mole-rats’ exceptional adaptations.

## Results

### Brain surgeries in mole-rats

The unique physiology and anatomy of *Fukomys* mole-rats require adaptations of brain surgery protocols typically employed for rats and mice (Dhawale et al., 2017; Okun et al., 2016). Each step, from managing anesthesia to addressing their specialized skull anatomy, demanded careful optimization. We describe the final protocol in detail in the Methods section. Below, we summarize the key challenges and the strategies developed to overcome them, ensuring both animal welfare and experimental success.

### Head muscles and skull thickness in mole-rats

To support surgical planning for implanting electrophysiology electrodes in the brain, we performed high-resolution *X-ray* micro-computed tomography (µ-CT) of iodine-stained heads of three mole-rat species (*Fukomys anselli, Fukomys darlingi, Heterocephalus glaber*) and the laboratory mouse (*Mus musculus*) to visualize the anatomy of the skull and overlying muscles (Figures S1 and S2). This revealed hypertrophied muscles in mole-rats, including thicker temporalis muscles that originate on the dorsal region of the skull (i.e., Os frontale and Os parietale) and extend from the midline suture down to the lower jaw. In stark contrast to mice, where the temporalis muscles are laterally positioned, these muscles cover large portions of the dorsal skull in mole-rats (Figure S1). These dorsal muscles had to be removed to gain access to the skull. To minimize pain and prevent the muscles from dehydration, we injected a sufficient volume of local anesthetic bupivacaine subcutaneously and constantly hydrated the muscles by flushing them with sterile 0.9% saline solution throughout the surgery. Surprisingly, the dorsal bones of the Ansell’s mole-rat skull are mostly thin (Figure S2, Table S1, down to 126 µm, n = 4 animals), but a mid-sagittal ridge that develops with age, protrudes up to 1780 µm in adults (mean ± SD, 1225 ± 206 µm, n = 4 animals, Figure S2). Due to the thin skull bones, screws to stabilize implants must be inserted with utmost care.

### Analgesia, anesthesia and surgery strategy

The implantation of recording electrodes requires appropriate anesthetic and analgesic measures. As the metabolism of mole-rats differs considerably from that of rats and mice (Gerhardt et al., 2023; Park et al., 2017) we first set out to establish stable surgical procedures. Here, we present data from 18 successful surgeries on mole-rats. We used isoflurane anesthesia because it allows precise control of depth and duration. Due to the adaptations to hypoxic environments, we used compressed air instead of pure oxygen as a carrier gas. A guinea-pig bite bar was attached to a custom-built nose cone to deliver the gas in a stereotaxic frame for mice.

During the surgery, it is important to keep the core body temperature of the animal stable. The body temperature of Ansell’s mole-rats (∼35°C, (Marhold & Nagel, 1995)) is lower than that of mice (37°C). The target body temperature on the heating pad was 34.9°C. To prevent dehydration during the surgery, the animals received a subcutaneous saline injection (2% of the animal’s body mass) at the beginning. After induction with 3% isoflurane, the animals reached a stable surgical depth anesthesia at concentrations ranging from 1.00 to 2.75% (Figure S3). This concentration had to be carefully monitored throughout the surgery, as the vital signs of mole-rats under anesthesia are considerably more variable than those of mice. For example, the breathing rate ranged between 10 to 80 breaths per minute (bpm). The average breathing rate decreased from 51 bpm to 32 bpm over 3 hours (Figure S3). Additionally, regular periods of not breathing occurred, a common characteristic of subterranean mole-rats, as they can hold their breath in hypoxic underground environments (Park et al., 2017). The duration of the anesthesia should not exceed 3 hours, after which the animals became more unstable. For this reason, we decided to split the surgical procedures into two steps, with a recovery period of 3-7 days between them (Figure S3). During the first surgery, the muscles were cleared, and the skull was cleaned, roughened, and coated with a thin base layer of dental composite. During the second surgery, the craniotomy was performed, and electrodes were inserted for acute recordings during the experiment sessions, whereas electrodes were cemented for chronic recordings.

### Post-operative care

Appropriate post-operative care was critical for the success of the surgeries. Most of the initial losses occurred during the recovery phase (80%), rather than during the surgery itself (20%). Across 45 surgeries on 33 mole-rats, intraoperative mortality was low (3 animals), with most losses occurring during the early postoperative recovery phase (12 animals). Following optimization of surgery and recovery conditions, 18 animals recovered successfully. We found that the most important factors in promoting animal survival were related to analgesia and how the animals were housed during post-surgical recovery.

### Analgesia

Mole-rats show greater sensitivity to the side effects of analgesics such as opioids (Kanui & Hole, 1990) and NSAIDs, which required adjustments of protocols. For example, we found that the standard dosage of 0.1 mg/kg of Buprenorphine sometimes resulted in a fatal shutdown of the gastrointestinal tract. Conversely, multiple doses of NSAIDs cause inflammation and hemorrhaging of the gastrointenstinal tract (McEvoy et al., 2021). We found that administering a single low dose of Buprenorphine (0.05 mg/kg) before surgery and daily single-dose administrations of the NSAID Carprofen (4 mg/kg) over three days post-surgery provided safe and sufficient analgesia.

### A KCC2 mutation that promotes seizures under normocapnic conditions

Mole-rats are adapted to live in environments with low oxygen and elevated carbon dioxide concentrations, particularly in their nest chambers, where they pile up to sleep (Vavrušková et al., 2022). These adaptations include ion channels that regulate the excitation-inhibition balance of the brain. Specifically, naked mole-rats (*Heterocephalus glaber*) and Damaraland mole-rats (*Fukomys damarensis*) were recently reported to possess a single amino acid substitution (R952H or R952C) in the neuronal potassium-chloride cotransporter 2 (KCC2) (Zions et al., 2020). The R952H mutation was also found in humans with febrile seizure disorder (Puskarjov et al., 2014). KCC2 participates in the maintenance of extracellular chloride ion concentrations for the efficient functioning of the brain’s inhibitory GABAergic system (Deeb et al., 2011; DeFazio et al., 2000). This function is likely compromised due to the mutation, but it is speculated that the inhibitory effect of increased CO_2_ concentrations in the underground tunnel system of mole-rats counterbalances this (Roper et al., 2001; Zions et al., 2020). The KKC2 mutation may therefore be of adaptive value as it saves energy in an underground environment, but under normocapnic conditions in a laboratory, it increases the likelihood of overexcitation, resulting in seizures (Zions et al., 2020). In our initial experiments, we noted that mole-rats kept under normal air CO_2_ concentrations after surgery, separated from the rest of their families to protect the wound and drives, often developed seizures.

Analysis of transcriptomic data from cortical tissue revealed that the R952C substitution in the KCC2 protein sequence is also present in our model species *Fukomys anselli* (Figure S4A), making it the third mole-rat species in which a mutation at this position has been reported. We suspected that this mutation, along with the resulting excitation-inhibition imbalance under normocapnic conditions, was responsible for the high seizure rate observed in our initial surgeries. To test this hypothesis, we performed CO_2_ measurements in the home and recovery cages (Figure S4B). These measurements revealed concentrations of 0.4 - 0.9% CO_2_ (mean ± SD: 0.58% ± 0.17%) in the nest boxes within the home cages, which was significantly higher than the 0.05 - 0.1% CO_2_ (0.07% ± 0.02%) in the cages that were initially used for recovery (One-Way ANOVA with Tukey’s post hoc multiple comparisons, F = 66.56, p < 0.0001, Figure S4B). Values similar to those found in our nest boxes have been reported for burrows of naked mole-rats (0.4% CO_2_) (Holtze et al., 2018) and *Fukomys damarensis* (mean 0.4% CO_2_, max 6% CO_2_) (Roper et al., 2001) in the wild. To increase the CO_2_ concentrations to those found in the nest boxes of their home cages, we started housing the experimental animals in minimally ventilated cages. The reduced ventilation in these cages caused CO_2_ to build up (0.14 - 0.68%, 0.35% ± 0.15%, One-Way ANOVA with Tukey’s post hoc multiple comparisons, F = 66.56, p < 0.0001) and efficiently reduced the occurrence of seizures. While this was a significant step towards the final protocol, there was yet another change to post-surgical care that proved significant: good company.

### Sociality

Mice and rats are mostly housed singly after implanting electrodes to minimize the risk of damage to the wound and the sensitive recording equipment (Kim et al., 2020). African mole-rats of the genus *Fukomys*, however, live in highly social family groups with one breeding pair and several generations of their offspring. We found it vital for the recovery to provide the company of a single family member of the same sex during the pre-surgical 2-week habituation period and the recovery period post-surgery, as this reduced the stress from isolation from the rest of the family group, and allowed huddling behavior which saves energy and increased CO_2_ levels (Okrouhlík et al., 2025; Vavrušková et al., 2022; Wiedenová et al., 2018). We placed a cage containing the two separated animals inside the family home cage to allow for vocal and olfactory communication with the rest of the family through holes in the lid of the cage. After the implant surgery, we found that adding the second animal to the cage for only a few hours a day under supervision was sufficient. This minimized the risk of the conspecific damaging the implant. The conspecific-cohousing time was gradually reduced until the implanted animal had fully recovered.

### Stereotaxic brain atlas of *Fukomys* mole-rats

The targeting of brain structures with the recording electrodes is achieved using stereotaxic coordinates taken from a brain atlas (Cecyn & Abrahao, 2023). Here, we present a stereotaxic brain atlas for the mole-rat *Fukomys anselli*, incorporating Nissl-stained coronal sections from two adult males. The atlas comprises 175 plates with 256 annotated brain regions and stereotaxic coordinates, based on Bregma and Lambda reference points (Figure S5). The annotation is based on anatomical comparisons with published atlases of the mole-rat *Fukomys anselli* (Dollas et al., 2019), mouse (Paxinos & Franklin, 2019), and rat (Paxinos & Watson, 2006). This novel resource enables the reliable targeting of brain regions during electrophysiological recordings and can serve as a reference for future mole-rat brain studies. It will be updated regularly as new anatomical and functional data become available. The current version of the atlas is available at https://doi.org/10.17617/3.UNDKRO.

### Electrophysiological recordings

The developed surgery protocol allowed us to perform electrophysiological recordings in awake mole-rats using tetrodes and Neuropixels (NPX) silicon probes. Below, we present findings from acute recordings in head-fixed animals followed by chronic recordings in freely moving *Fukomys* mole-rats.

### Repeated acute Neuropixels recordings in head-fixed mole-rats

Neuropixels probes allow the simultaneous recording of hundreds of neurons across multiple brain regions (Jun et al., 2017). We used these probes in awake head-fixed *Fukomys* mole-rats, targeting the medial visual cortex (V2M), the retrosplenial cortex (RSC), and the superior colliculus (SC). Here, we report data from acute recordings in response to auditory and visual stimulation in two head-fixed awake mole-rats running on a moving disk (Figure 2A). We used Neuropixels 1.0 probes that were lowered into the brain at the predetermined insertion locations based on the mole-rat stereotaxic brain atlas (∼5.0 mm posterior to bregma, ∼1.5 mm lateral to the midline). Following the electrophysiological experiments, we reconstructed the electrode tracks and matched them to the brain atlas (Figure 2B). We employed a two-step histological procedure to achieve this: 1) whole brain tissue clearing (modified iDISCO^+^ protocol (Renier et al., 2016), Table S2) followed by fluorescence light-sheet microscopy to visualize the probe tracks in 3D (Video S1) and 2) a reverse clearing protocol (reDISCO, Table S3) followed by vibratome sectioning, DAPI staining, and fluorescent microscopy to reveal the brain areas in more detail (Figure 4A).

**Figure 2:**
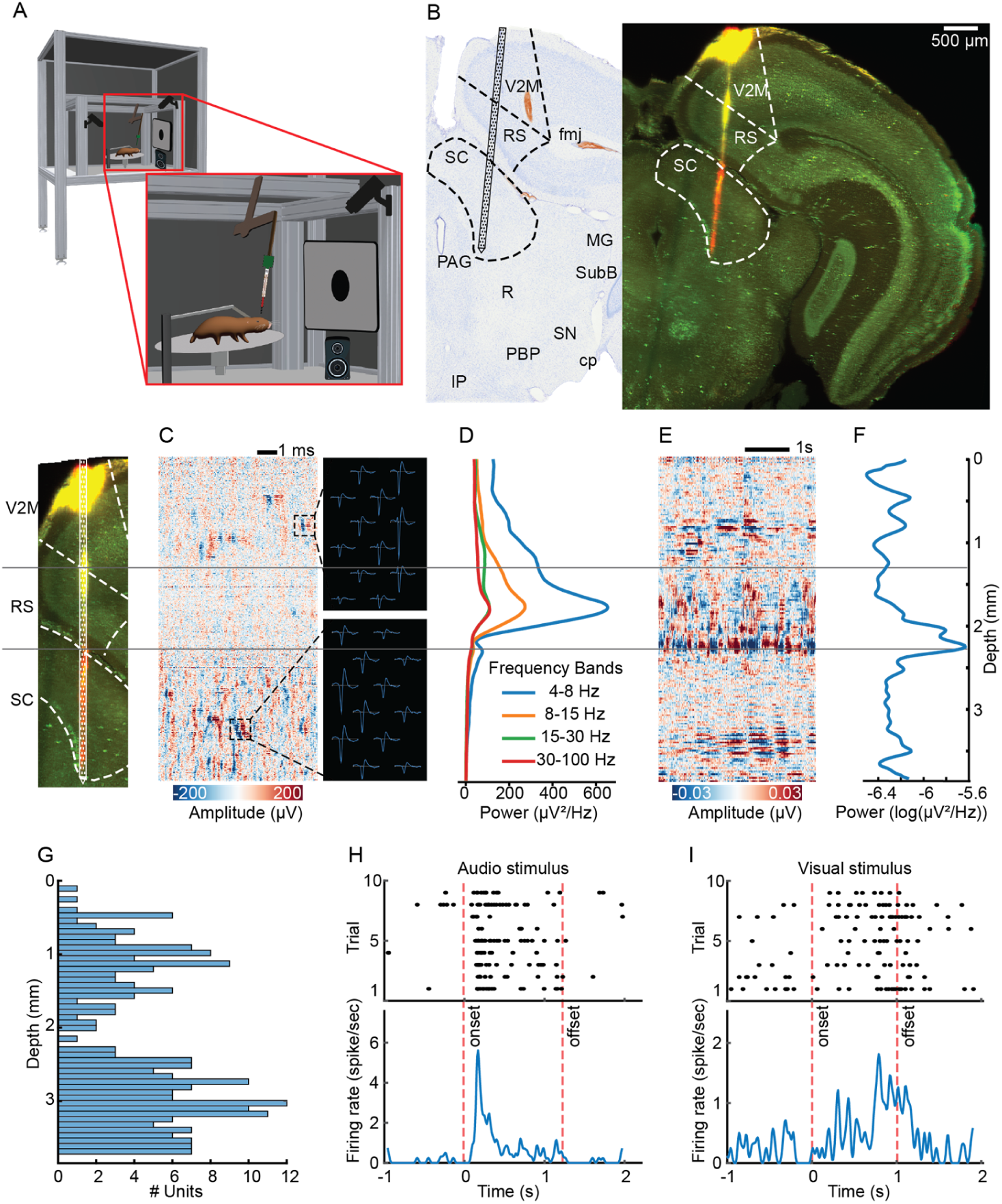
Acute Neuropixels recordings in the superior colliculus in a head-fixed *Fukomys* mole-rat reveal units responding to auditory or visual stimulation. **A.** The setup with a head-fixed African mole-rat running on a disk. Visual and auditory stimuli are presented via a projector screen and loudspeaker in front of the animal, respectively. **B.** Histological reconstruction (right, virtual section from light sheet microscopy data) of the probe track labelled with CM-DiI (red) in cleared tissue and the corresponding image (left, Plate 101) from the <u>stereotaxic mole-rat brain atlas</u>. The borders of recorded brain regions are shown with dashed lines. **C.** Example snippet of preprocessed high-frequency data showing increased spike densities in cortex and superior colliculus. **D.** LFP power in different frequency bands, computed by Welch’s method in a 1-s interval, averaged over ten intervals. **E.** Example snippet of current source density, computed as the second-order spatial derivative of broadband LFP data (0.5-25 Hz). **F.** Current source density power. **G.** Distribution of recorded units along the electrode depth in one example recording session. **H. and I.** Raster plots and peristimulus time histograms of example auditory and visual (respectively) responsive cells recorded from SC. The dashed lines indicate the stimulus onset and offset.

Several measures of neural activity were used to confirm that the electrophysiological recordings aligned with the probe’s anatomical position identified in histological reconstructions. First, high-pass filtered, common reference averaged data (Figure 2C) showed different activity patterns along the probe, revealing the laminar structure of the targeted area. Second, local field potential (LFP) oscillations have different profiles across brain regions (Buzsáki et al., 2013). Therefore, we used the frequency of the recorded LFP as an additional electrophysiological measure to improve/validate the probe location in the brain. The amplitude of distinct frequency ranges reflected the border between cortex and subcortical areas (e.g., SC, Figure 2D). Third, current source density (CSD) analysis, which transforms LFP signals into locations of current sources and sinks, has been used to reveal a laminar distribution of neuronal activities in different brain structures and animal models (Kaur et al., 2005; Niell & Stryker, 2008; Swadlow et al., 2002). Here, we used CSD analysis to infer the laminar profile of LFP traces and to further improve/validate the alignment of electrophysiological signals with histology. As shown in the example recording, the CSD traces and their power capture laminar structures, highlighting again the transition between the cortex and the SC (Figure 2E and F).

In the next step, we isolated single unit activity using automated spike sorting methods (Kilosort 4) (Pachitariu et al., 2024) with manual curation (Phy 2) (Rossant et al., 2016). Overall, 1,210 units were isolated from the cortex, SC, and PAG in two animals. The distribution of the recorded neurons along the probe depth matched with the boundaries of anatomical structures along the probe (note the silent region between the RSC and the SC, Figure 2G). We found neurons in the SC that responded to auditory (Figure 2H) or visual (Figure 2I) stimulation. In summary, these results demonstrate that our method enables acute neuronal recordings with Neuropixels in head-fixed mole-rats to measure LFP and single-unit responses to sensory stimulation, and reconstruct the anatomical location of these recordings.

### Chronic electrophysiology recordings in freely moving mole-rats

Chronic electrophysiology recordings enable the study of long-term dynamics of neural activity in behaving animals (Guan et al., 2019; Saeedi et al., 2024; Steinmetz et al., 2021). Repeated recordings from the same neurons allow the study of neural plasticity, adaptation, and circuit-level changes in response to environmental stimuli and behavioral adaptations, such as cooperative foraging, dominance hierarchies, and responses to environmental threats (Steinmetz et al., 2021). Here, we established chronic electrophysiology recording methods in *Fukomys* mole-rats using wire electrodes (tetrodes), which permitted stable recordings from a target brain area well as Neuropixels probes, which enabled simultaneous recordings from multiple brain areas.

### Chronic tetrode recordings reveal low frequency theta rhythms in the hippocampus of freely moving *Fukomys* mole-rats

The hippocampus is involved in cognitive processes such as memory and navigation in which oscillatory activities play a crucial role (Chrastil et al., 2022; Colgin, 2016). One of the most prominent oscillations in the mammalian hippocampus is the theta rhythm, a low-frequency (< 20 Hz) oscillation that supports neural synchronization and temporal spike phase coding (Buzsáki et al., 2003; Vanderwolf, 1969). We recorded local field potential (LFP) and stable single-unit spike data (Figure 3C) using tetrodes positioned in the hippocampal CA1 region of two female mole-rats (age 22 months and 13 months) exploring a narrow circular track (Figure 3A). We subsequently computed spectrograms (Figure 3G) and the power spectral density (Figure 3D) of the low-pass filtered LFP signal to analyze the theta rhythm. This revealed a low-frequency band with a prominent peak between 4 and 5 Hz (Figure 3D). To determine the average frequency of this suspected theta band, we computed the frequency range of the peak at full width at half maximum across recording sessions in two animals. The results showed a theta band ranging from 3 to 6 Hz (n = 32 recording sessions, mean ± SD, lower bound: 3.25 ± 0.55 Hz, upper bound: 5.97 ± 0.41 Hz) (Figure 3F). This oscillation in mole-rats is lower than the theta frequency band observed in other rodent species, such as rats (Vanderwolf, 1969) and mice (Buzsáki et al., 2003) (6 to 12 Hz). In summary, our tetrode recordings revealed a theta oscillation in the hippocampal LFP of mole-rats that is low for a rodent.

**Figure 3:**
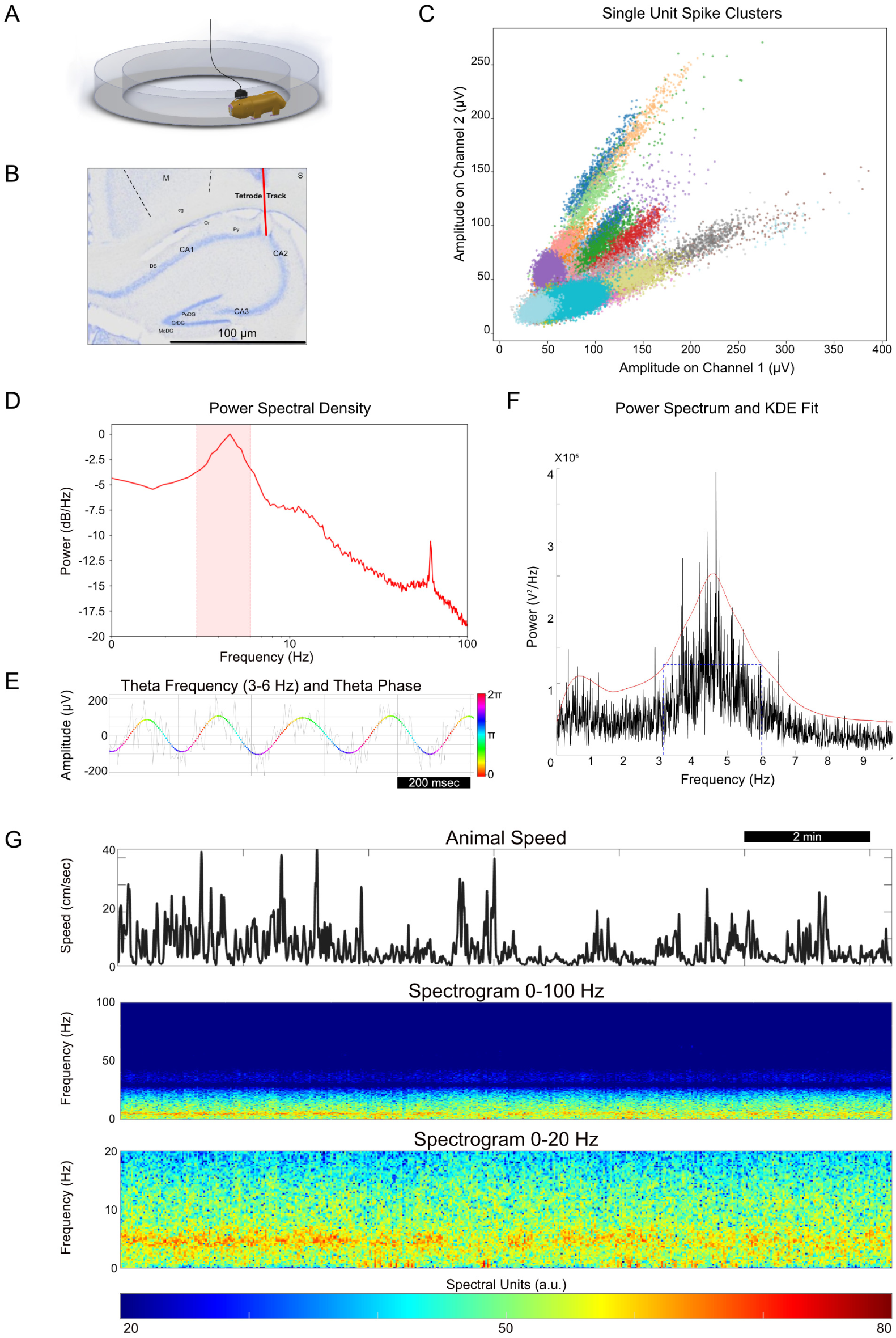
Tetrode recordings from the hippocampus of freely moving Ansell’s mole-rats reveal a low-frequency theta band. **A**. Recording setup. Circular track explored by the animal during recording sessions. **B.** Histology. Brain section with electrolytic lesion at the end of the tetrode track in the hippocampus CA1 area (bottom). **C**. Single-unit spike clusters. Illustrative clusters of single-unit data recorded simultaneously from two tetrodes in the CA1 area. **D.** Power spectral density (PSD). PSD plot from a single recording session, highlighting the prominent 3 to 6 Hz theta frequency band. **E.** Theta trace. Raw LFP trace (grey) (1 second window) from the same session and its theta filtered trace (colored), showing the corresponding phase in the range 0 to 2π radians. **F.** Theta range analysis. Power spectrum of an example session with a fitted kernel density estimate (KDE fit) and its full width at half maximum, which was used to determine the frequency range of mole-rat hippocampal theta (right). **G.** Speed and LFP spectrogram. Example speed plot (top) of a single recording session (12 min) with corresponding LFP spectrogram in the range of 1 to 100 Hz (middle) and 1 to 20 Hz, showing a prominent theta band between 3 and 6 Hz in the (bottom).

### Chronic Neuropixels recordings in freely moving mole-rats

Although tetrodes yield stable chronic recordings, they are limited in the number of neurons and brain areas that can be recorded from simultaneously. In line with the 3R principles to reduce the number of animals used in experiments, however, it is crucial to maximize the amount of data obtained from each animal. We were therefore motivated to increase the neuron yield and set out to establish chronic high-density electrophysiological recordings in freely moving mole-rats using Neuropixels 2.0 probes. These probes have 5,120 electrode sites distributed across four shanks, each 10 mm long, enabling simultaneous recordings from 384 channels (one bank) (Steinmetz et al., 2021).

We recorded from two adult Ansell’s mole-rats (one male, one female) using a customized version of the recently developed Apollo implant (Bimbard et al., 2025). The probes were implanted to record activity in the retrosplenial cortex (RSC), hippocampus, and thalamus. Reconstruction of the probe track confirmed that the probes targeted these areas (Figure 4 and Figure S6). We were able to record high-quality electrophysiological signals for more than four weeks after implantation (34 days in animal 1, 30 days in animal 2, example traces in Figure 4C). Each day of the experiment, we recorded neuronal activity from either the upper or lower 384 channels of one shank. Overall, the number of units recorded by the lower banks of the probe (deeper structures, hippocampus, and thalamus) was higher than the units recorded by the upper banks (mainly RSC), but the number of units did not decline over the course of three weeks (Figure 4D). Quality metrics of the units, such as firing rate, signal-to-noise ratio, and amplitude, also remained consistent throughout the entire recording period (Figure 4E-G). The stable low-noise levels and high spike amplitudes of the recorded units demonstrated that NPX2.0 probes can be used for long-term studies in mole-rats. In summary, we have set up a chronic recording system that enables high-yield single-unit recordings in freely moving mole-rats, enabling studies of brain activity during natural behaviors.

**Figure 4:**
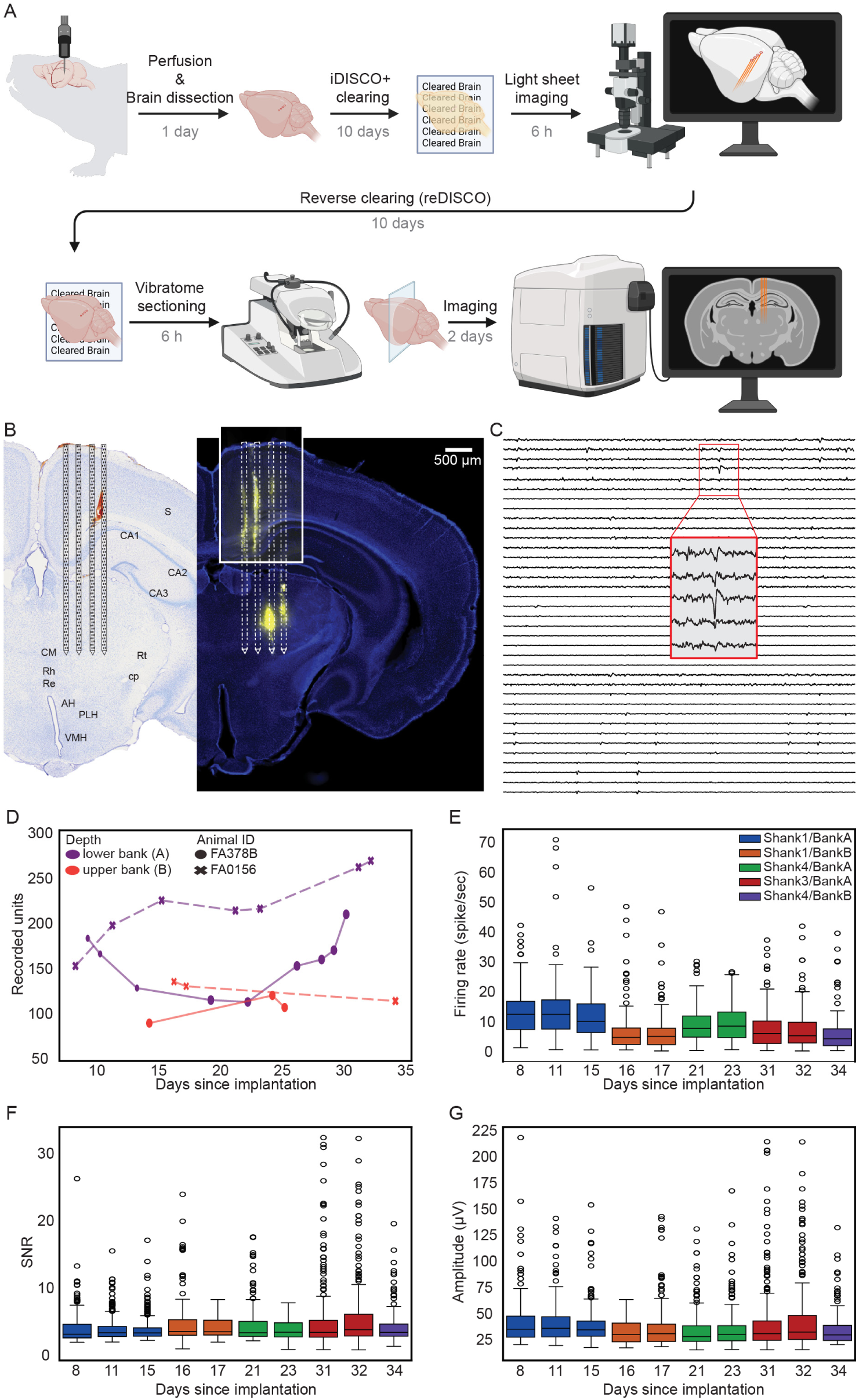
Chronic electrophysiology in freely moving Ansell’s mole-rats using Neuropixels 2.0 probes. **A.** Workflow and approximate timeline of the histological reconstruction of the recording sites. The electrode tracks were determined in a two-step procedure. First, the brains were cleared using a modified iDISCO^+^ protocol, followed by light sheet microscopy to reconstruct the electrode tracks in three dimensions. After imaging, the brains were rehydrated, sectioned at a vibratome, counterstained with DAPI and a fluorescent Nissl stain and imaged at a fluorescent slide scanner. Created with BioRender.com. **B.** Histological reconstruction of the four NPX 2.0 probe tracks with CM-DiI (yellow) in a 100 µm brain section alongside the corresponding plate from the stereotaxic mole-rat brain atlas. **C.** Example snippet of preprocessed high-frequency (AP) data from a subset of channels showing the spiking activity. **D.** Number of recorded units on different shanks of NPX 2.0 from two Ansell’s mole-rats over the course of several weeks. No decline in neuron numbers was observed. **E.** Firing rates of recorded units from one mole-rat over the course of one month. Different colors indicate the shank and bank in the NPX 2.0 probe used for data collection. **F.** Signal-to-noise ratio (SNR) and **G.** amplitude (µV) of the recorded units, respectively. The color code is as in E.

## Discussion

In this paper, we describe how we established electrophysiological recordings in subterranean *Fukomys* mole-rats. We summarize the key elements of standard protocols for intracranial surgeries in rodents that had to be modified to accommodate these animals’ adaptations to their unique subterranean habitat, and we present findings from acute and chronic recordings.

The first challenge was the anatomy of the mole-rat’s head. Most bathyergid mole-rats are chisel-tooth diggers, requiring long, sharp incisors and strong jaws to tunnel their burrows and manage their diet of hard roots and tubers (Cox et al., 2020; Van Daele et al., 2009). The strength required for this digging and chewing depends on the bone and muscle structures that contribute to their strong bite force. It is mediated by a large temporalis muscle, which is not only thick, but, in contrast to mice, also extends dorsally up to the midsagittal ridge. The thick muscles complicate surgeries on the dorsal part of the head and the careful removal of these muscles significantly increases the length of the surgery. Second, we had to develop protocols for controlled anesthesia and analgesia. Due to their low metabolism and body temperature, mole-rats have been reported to require significantly lower doses (∼10 times less) of injectable anesthetics (Ketamine/Xylazine) than other rodents for minimally invasive surgical interventions (Garcia Montero et al., 2015). However, protocols for long and invasive brain surgeries had not yet been established. The challenge was to identify the lowest tolerable dose while maintaining sufficient anesthetic depth and analgesic efficacy throughout the lengthy procedure. We found that isoflurane provides safe anesthesia, and that it is particularly important not to overdose mole-rats with opioids and NSAIDs, as their gastrointestinal tracts are overly sensitive to the known side effects of these drugs, such as constipation and bleeding. The cause of the potentiation of these potentially fatal side effects is unknown. We speculate that the hypoxic conditions in the habitat of the mole-rats, which are known to downregulate the expression of key enzymes in the metabolism of drugs in the liver and gastrointestinal tract (e.g. cytochrome p450) might play a role (Wang et al., 2025; Zhou et al., 2018).

Once stable surgical procedures had been established, the conditions during the first two recovery days proved crucial to survival. Mole-rats possess numerous physiological adaptations to the hypoxic and hypercapnic underground environment, including a reduced resting metabolic rate (Schielke et al., 2017; Zelová et al., 2007) and an increased affinity of hemoglobin for oxygen (van Aardt et al., 2007; Weber et al., 2017). The brain, as a metabolically expensive tissue, can be expected to have evolved specific adaptations to cope with the energy constraints imposed by altered oxygen and carbon dioxide levels. In our study species, *Fukomys anselli*, we identified a mutation in the neuronal potassium-chloride cotransporter 2 (*KCC2*) gene. A similar mutation, previously reported in other mole-rat species, including the naked mole-rat (Zions et al., 2020), presumably reduces GABAergic inhibition in the brain. It may represent an adaptation to subterranean life, where elevated carbon dioxide levels naturally inhibit neural activity. By reducing the energetic cost of active inhibition via GABAergic signaling, the mutation could contribute to energy conservation in a hypercapnic burrow environment (Zions et al., 2020). Conversely, under normocapnic laboratory conditions, particularly following invasive brain surgery, this mutation appears to increase the risk of seizures during the postoperative recovery period. We show that this vulnerability can be mitigated by housing the animals under elevated CO_2_ conditions in minimally ventilated cages after surgery. Furthermore, we found that the presence of family members during the early recovery phase is beneficial. Social African mole-rats live in multigenerational family groups, and social contact is important for their well-being. They show characteristics of heterothermia and rely on behaviors such as huddling in their nest to maintain their body temperature at 33 - 35°C (Marhold & Nagel, 1995; Okrouhlík et al., 2025; Vavrušková et al., 2022; Wiedenová et al., 2018). In addition, multiple animals cause faster and greater CO_2_ build-up in the cage, which, as mentioned above, helps to prevent seizures. Although housing implanted animals with their family members supports recovery by promoting natural social behaviors and a stable microenvironment, it also increases the risk of damage to the implanted recording device. In our experiments, animals were paired with one same-sex conspecific from up to two weeks prior to the first implant surgery, throughout the entire recovery period following the first surgery, and during the initial post-operative days after the second surgery. To minimize the risk of damage to the drives, we housed the implanted animal with a single, same-sex family member only, progressively reducing the daily time of social contact until the implanted animal was housed alone after completing its recovery from the second surgery.

Using the established surgery and recovery protocols, we demonstrated the ability to record from the brains of mole-rats in both acute head-fixed and chronic freely-moving settings using two recording techniques. We developed a novel stereotaxic framework for the Ansell’s mole-rat brain to guide the electrode placement. Acute Neuropixels recordings allowed to capture neuronal activity across cortical and subcortical target regions. The precise histological alignment was supported by the agreement between the electrophysiological activity and the histological boundaries. For example, we isolated signals from putative single neurons in the superior colliculus and recorded their response to auditory and visual stimuli.

Next, we used tetrodes to record from the dorsal hippocampus of freely moving Ansell’s mole-rats. As with all other rodents studied so far, we found that mole-rats show a strong rhythmic oscillation in the theta frequency range, supporting the correct positioning of the tetrodes in the CA area. Interestingly, theta in mole-rats is in the range of 3 to 6 Hz, shifted towards lower frequencies compared to other rodent species. Rats (Vanderwolf, 1969) and mice (Buzsáki et al., 2003) have theta frequency bands around 6 to 12 Hz and similar ranges have been reported for other rodents, including guinea pigs (Sainsbury & Montoya, 1984) (5-10 Hz) and Mongolian gerbils (Whishaw, 1972) (7-10 Hz). This lower frequency range observed in mole-rats fits better with non-rodent mammal species such as rabbits (Nokia et al., 2008) (3-8 Hz), ferrets (Dunn et al., 2022) (4-7 Hz), sheep (Perentos et al., 2022) (4-10 Hz), marmosets (Piza et al., 2024) (4-10 Hz) and macaques (Abbaspoor et al., 2023) (3-10 Hz).

Finally, we demonstrated that the neuronal yield from chronic recordings can be significantly scaled up using Neuropixels 2.0 probes, enabling daily recordings from hundreds of neurons under freely moving conditions over several weeks. We also present a protocol for reconstructing the probe tract in 3D using tissue clearing and light sheet microscopy followed by a more detailed investigation of the recording sites on brain sections after reverse tissue clearing, supported by the novel stereotaxic brain atlas.

In summary, we established electrophysiological recording protocols in *Fukomys* mole-rats, mammals uniquely adapted to life in a subterranean environment. Social African mole-rats, including the naked mole-rat, are valuable model organisms for longevity studies, cancer research, and hypoxia tolerance. Furthermore, they also serve as excellent model organisms for studying the enigmatic magnetic sense and complex social behaviors. We encourage their use in neuroscience and hope that the protocols provided here offer a foundational framework for a wider uptake of these uncommon models to investigate mammalian brain function in the context of extreme physiological and behavioral adaptations.

## Acknowledgments

We acknowledge funding from the Max Planck Society and the European Research Council (ERC StG No 948728 to EPM). Runita Shirdhankar received a doctoral scholarship from the DAAD and is supported by the International Max Planck Research School (IMPRS) for Brain and Behavior. We are indebted to the animal unit and core facilities at the MPINB.

## Author contributions

Conceptualization, Methodology, Validation, Formal Analysis, Investigation, Data Curation, Writing (Original Draft), Writing (Review and Editing), Visualization, R.N.S.; Conceptualization, Methodology, Validation, Formal Analysis, Investigation, Data Curation, Writing (Original Draft), Writing (Review and Editing), Visualization, A.S.; Methodology, Formal Analysis, Investigation, Data Curation, Writing (Review and Editing), G.E.F.; Investigation, Data Curation, Writing (Original Draft), Writing (Review and Editing), Visualization, L.M.; Formal Analysis, Data Curation, Writing (Review and Editing), P.-Y.J.; Investigation, L.M.W.; Investigation, S.W.-K.; Investigation, W.B.; Investigation, E.H.; Resources, Writing (Review and Editing), Supervision, S.B.; Conceptualization, Methodology, Validation, Formal Analysis, Investigation, Resources, Data Curation, Writing (Original Draft), Writing (Review and Editing), Visualization, Supervision, Project Administration, Funding Acquisition, E.P.M.

## Ethics

All animal experiments were performed in accordance with relevant institutional and national guidelines and regulations. All experiments were approved by the Landesamt für Verbraucherschutz und Ernährung (LAVE) NRW, Germany and conducted in accordance with the European Union Directive 2010/63/EU on the protection of animals used for scientific purposes.

## Declaration of generative AI in scientific writing

During the preparation of this work, the authors used ChatGPT in order to improve the readability and language of the manuscript. After using this tool/service, the authors reviewed and edited the content as needed and take full responsibility for the content of the published article.

## Data and code availability

All code is available at https://github.com/Malkemperlab/Molerat-Ephys

Raw data will be provided by the corresponding author on reasonable request.

## Methods

### Key resources table

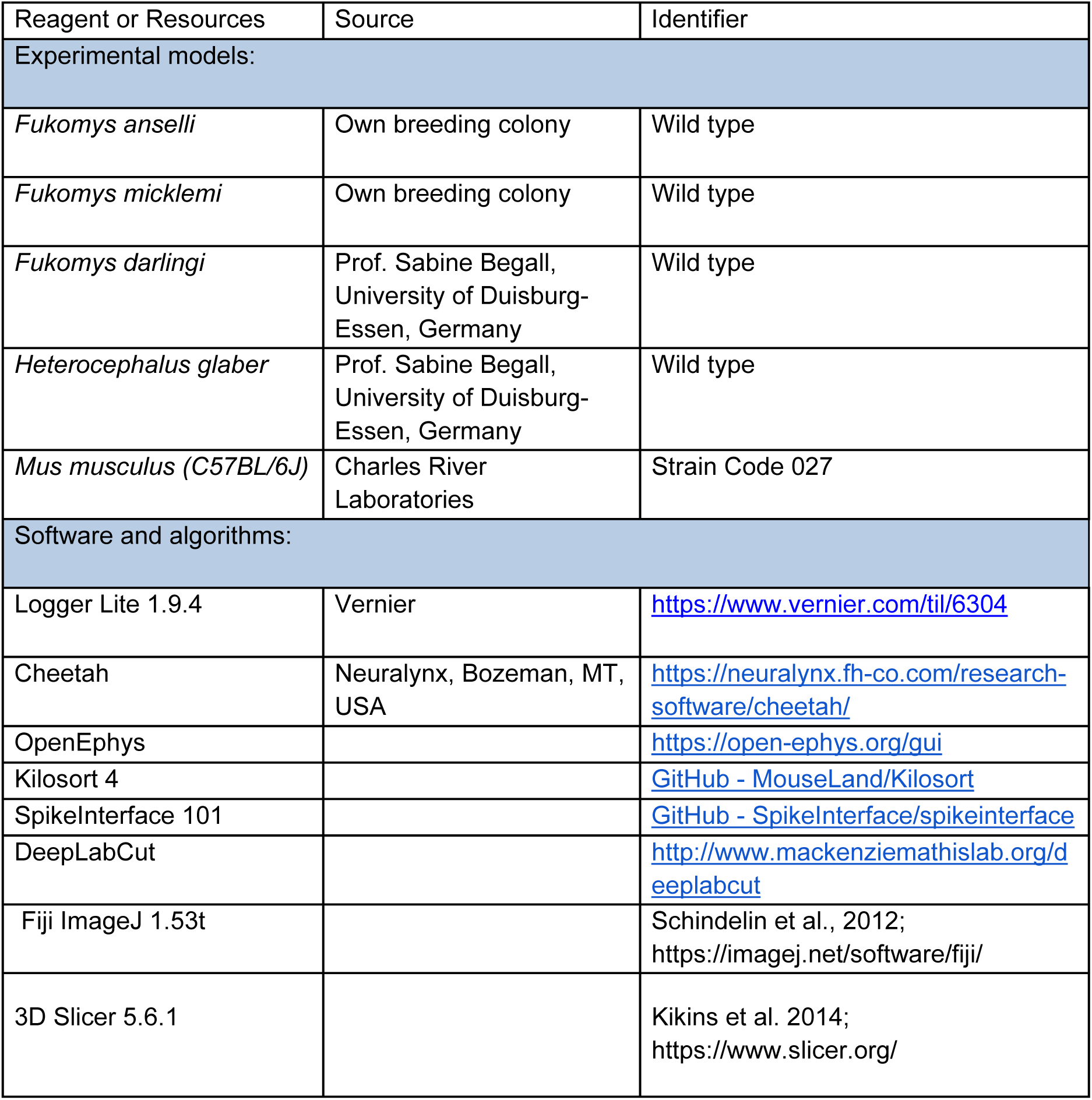
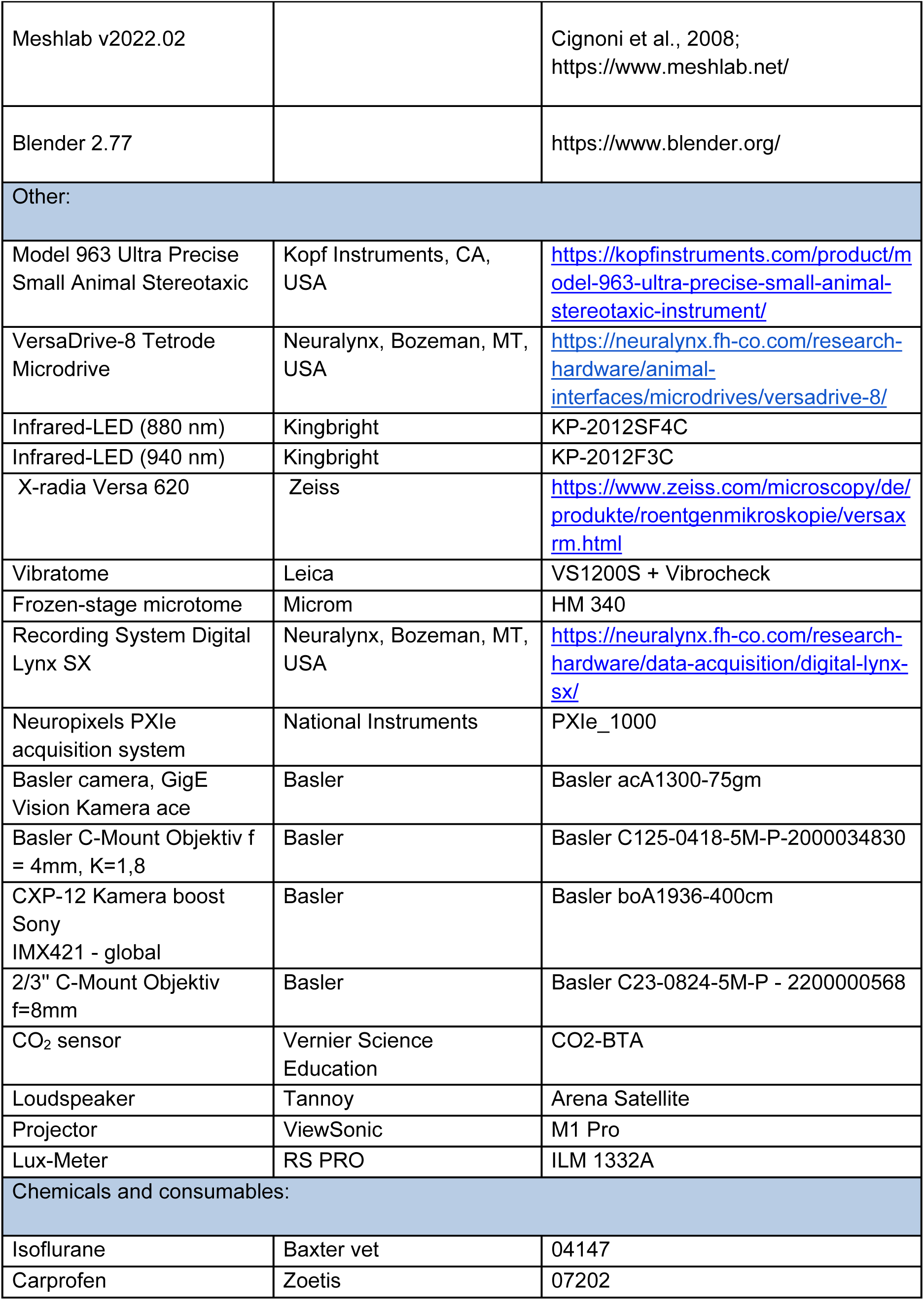
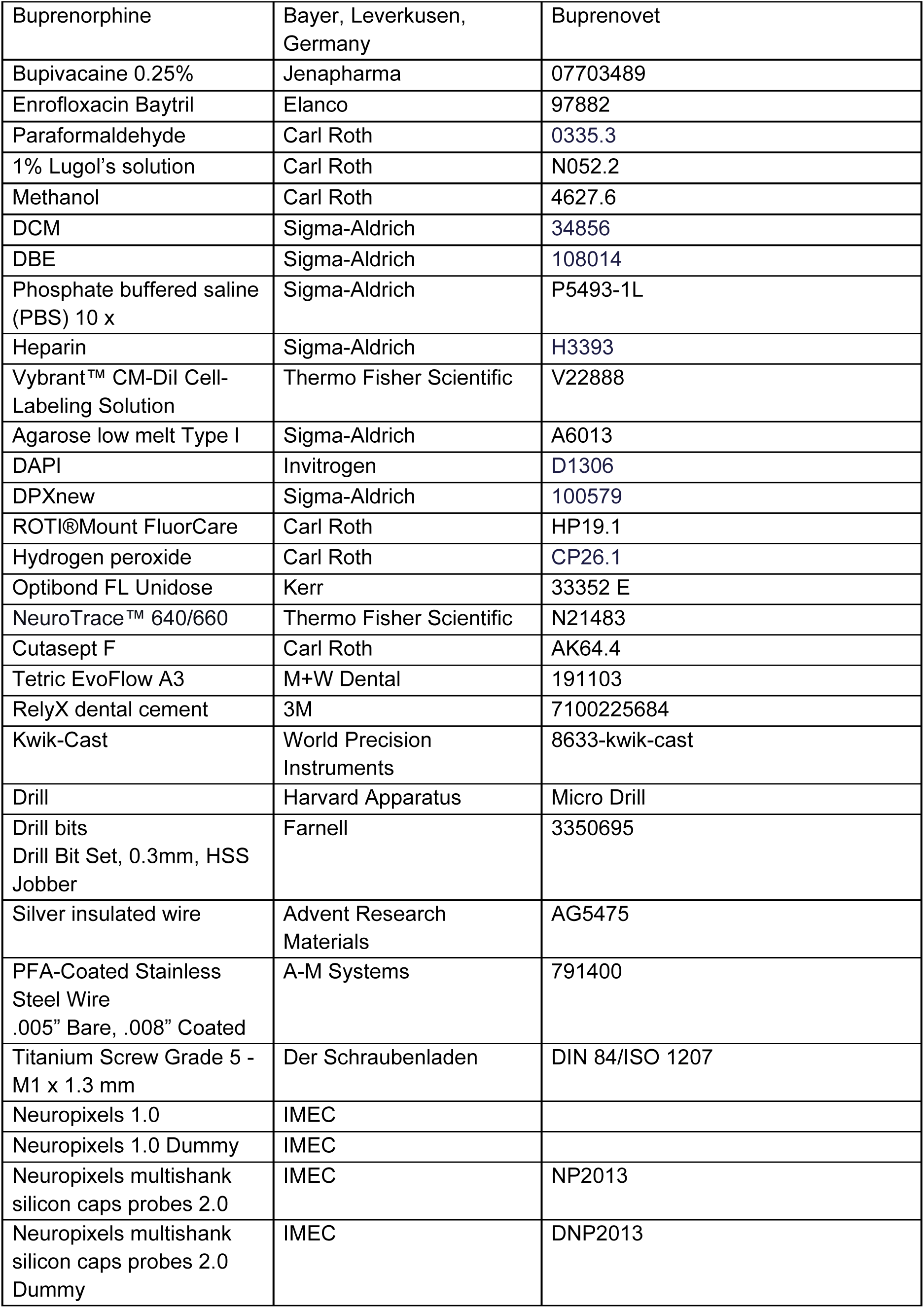
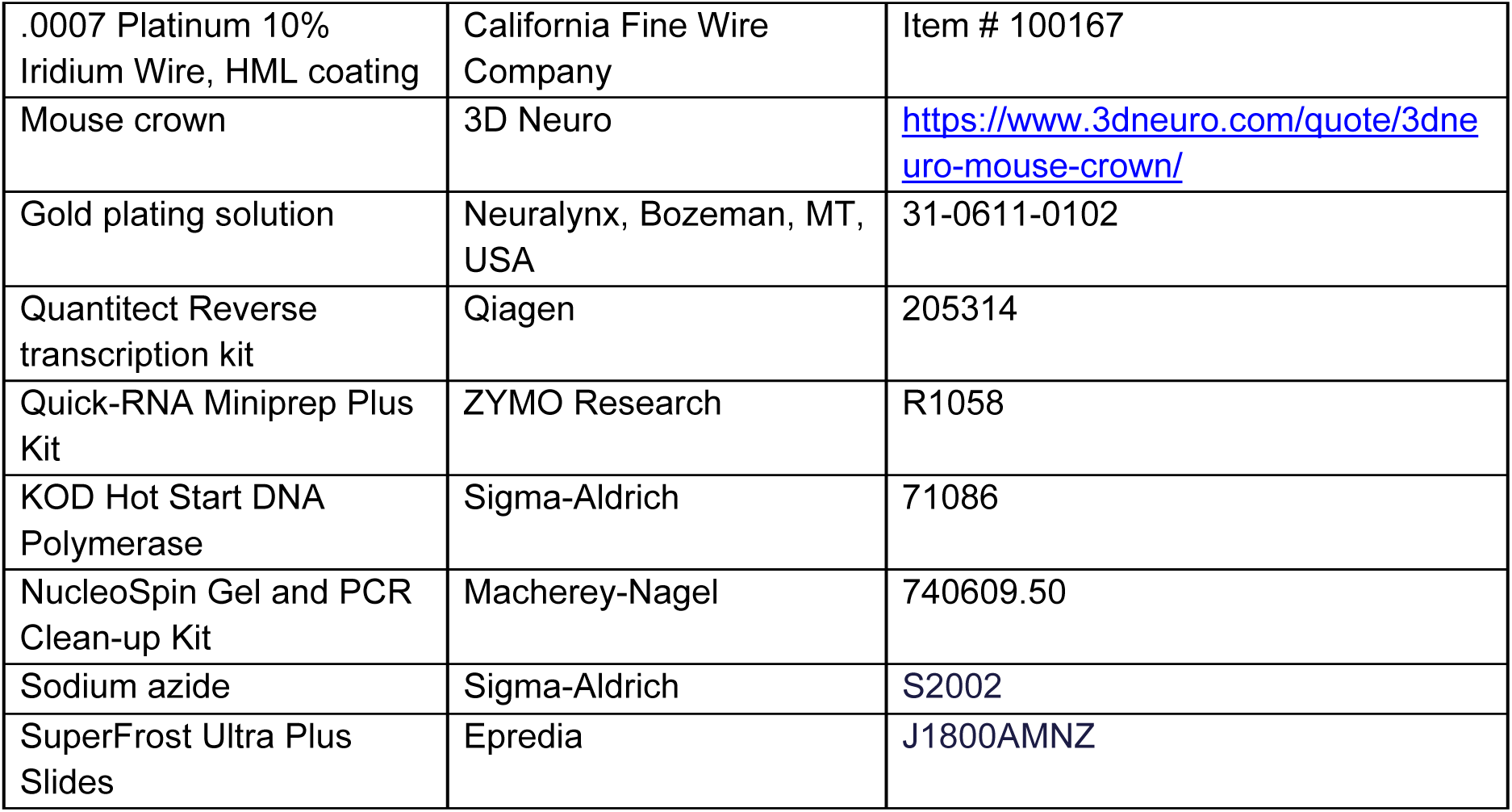

### Study organisms

Electrophysiological recordings were performed on two African mole-rat (Bathyergidae) species which according to recent phylogenetic analyses (Šumbera, pers. comm.) can be considered a single species: The Ansell’s mole rat *Fukomys anselli* and the Micklem’s mole-rat *Fukomys micklemi*, both included in the genus *Cryptomys* in older studies (Kocks et al., 2006). Two female Micklem’s mole-rats (age: 12 and 34 months) were used for head-fixed NPX recordings. Two female Ansell’s mole-rats, (age: 22 months and 13 months), were used for tetrode recordings in freely moving animals. Two freely moving Ansell’s mole-rats (age: female 29 months old and one male 19 months) were used for NPX recordings in freely moving animals.

### Surgical procedure and animal housing

To implant the electrophysiological recording apparatus, we performed intracranial surgeries on mole-rats in two stages, one week apart. This two-step approach minimized the duration of anesthesia and significantly reduced complications, which allowed for better recovery. All surgeries were performed in a high-precision stereotaxic apparatus (Kopf Instruments Model 963).

#### Animal housing

All mole-rats used in this study were bred in our own breeding colonies in a custom-made cage system. They were fed ad libitum with fresh vegetables including potatoes, carrots, cucumber and lettuce, supplemented with dry food pellets for guinea pigs (Altromin 3013). Mole-rats live in colonies typically consisting of a single breeding pair and their offspring, and therefore, social contact is essential for their well-being. However, housing implanted animals with their family poses significant risks, including potential damage to the implanted recording device or detachment of the implant from the skull. To reduce these risks, implanted animals were housed in a separate box (length x width x height: 56 cm x 39 cm x 21 cm) made from red perspex that was placed inside the home cage. A conspecific from the family was housed with the experimental animal from the first day of the isolation habituation period, which began two weeks before the first implant surgery. They were also co-housed during the recovery period following the first surgery. After the second surgery, the time spent with the conspecific was gradually reduced, and the implanted animals were housed alone after full recovery. This housing strategy balanced the social and physiological needs of the mole-rats while ensuring the integrity of the recording devices throughout the experiments.

#### Anesthesia and preoperative preparation

Buprenorphine (0.05 mg/kg) was administered subcutaneously half an hour before the surgeries to provide analgesia. Anesthesia was induced with 3% isoflurane in an induction box and maintained at 1−2.75% via a custom-made nose-cone during surgery. The scalp was shaved and sterilized with Cutasept F (Bode Chemie GmbH) before making incisions, and bupivacaine (0.2 - 0.4 ml of 0.25% solution depending on the animal’s head size) was injected subcutaneously under the scalp to provide local analgesia.

#### First surgery

In the first surgery, an incision was made along the dorsal region of the head to expose the temporalis muscle, which was carefully dissected and pushed to the sides to reveal the skull. The periosteum and connective tissue on the skull were thoroughly cleaned to ensure a smooth surface for implant fixation. Hydrogen peroxide (4% in sterile Milli-Q) was used to clean any residual tissue, followed by a wash with sterile saline to prepare the skull surface. For head-fixed recording experiments, a lightweight head plate (RODIN IECL-S Acute Implant, Cambridge NeuroTech) was installed onto the skull using an adhesive primer (OptiBond FL, Kerr Dental) and dental composite (Tetric EvoFlow, Ivoclar Vivadent). For freely moving experiments, a base layer of adhesive primer was applied to the cleaned skull, and a well was made using dental composite. The well was covered with Kwik-Cast (WPI), the skin was sutured, and animals were allowed to recover for at least 5 days.

#### Second surgery

After full recovery from the first surgery, mole-rats were anesthetized and placed in the stereotaxic frame as described above. The Kwik-Cast cover was removed to expose the skull. The craniotomy holes were drilled over target brain regions with a micro drill and carbide drill bits (Harvard apparatus) to allow electrode access.

For acute Neuropixels recordings, a silver wire (ADVENT, AG5475 Silver Insulated Wire) was implanted between the dura and the skull over the frontal lobe to serve as ground and reference and was fixed with dental composite to the head plate and skull. The craniotomy site was covered with Kwik-Cast.

For chronic Neuropixels recordings, a duratomy was performed with a 30G needle to partially remove the dura at the craniotomy site to facilitate the electrode insertion. A Neuropixels 2.0 probe with Apollo Implant assembly (Bimbard et al., 2025) was slowly lowered into the brain up to a depth of 5.5 mm from the surface. The implant was then fixed to the skull using dental composite. A silver wire, as ground and reference wire, was placed between the dura and the skull and fixed with dental composite as described above. A flexible 3D-printed cage (mouse crown, 3Dneuro) covered with copper mesh was installed around the implant to protect the probe.

For freely moving experiments with tetrodes, two holes were drilled into the skull for ground screw placement: one over the olfactory bulb and another over the cerebellum on the contralateral side of the skull. Screws (Titanium Screw Grade 5 - M1 x 1.3 mm) were placed into the ground holes. The screws were soldered to stainless steel wire (A-M Systems, 791400 PFA-Coated Stainless Steel Wire), which was then connected to the ground on the tetrode microdrive using a Mill-Max connector. In addition, three bone screws were secured into the skull to increase the stability of the implanted device. The recording device (see details below) was placed over the craniotomy, and the ground wires were inserted and secured using dental composite. After the second surgery the animals were allowed to recover for at least 3 days (head-fixed recordings) or 7 days (chronic recordings).

#### Postoperative analgesia

After each surgery, the animals received a daily subcutaneous dose of 4 mg/kg body mass Carprofen as analgesics, administered at least once and on the following days if required.

#### Head-fixed Neuropixels recordings

After recovery from the second surgery, mole-rats used for head-fixed experiments were habituated to the setup and head fixation on a spinning disc for three consecutive days (0.5 h/day) (Figure 2A). The animal was attached to the head-fixation system and a Neuropixels 1.0 probe connected to a hydraulic micromanipulator system (Narishige, MO-10) was slowly lowered into the brain (∼200 μm/min). After the probe was in place, the data acquisition was started, and the electrophysiology signals in response to sensory stimulation were collected using the Neuropixels data-acquisition system (Durand et al., 2023). The data from the Neuropixels probe was sampled at 30 kHz for action potential (AP) and at 2.5 kHz for local field potential (LFP) signals and saved for further analysis using the OpenEphys GUI (Siegle et al., 2017). At the end of the experiment, the probe was slowly retracted, the skull was covered using Kwik-Cast, and the animal received a food reward and was brought back to the home cage. Recordings continued on consecutive days as long as neuronal isolation remained of high quality.

#### Auditory stimulation

We generated auditory stimuli digitally at a sampling rate of 44,100 Hz as follows and presented them over 1.2 seconds. Frequency-modulated tone: A pure tone with an intensity of 70 dB SPL linearly modulated in frequency from 200 Hz to 1,400 Hz across the stimulus duration. The stimuli were presented through a Tannoy Arena Satellite speaker placed ∼ 40 cm in front of the animal.

#### Visual stimulation

Two types of visual stimuli were presented using a mini-projector (Viewsonic M1 Pro) and screen (30 cm x 40 cm) placed 38 cm in front of the animal. All stimuli consisted of a single dot displayed on a uniform gray background using Psychtoolbox (MATLAB). Bright Dot (Condition 1): A white circular dot was presented centrally and increased in size from 4° to 40° of visual angle, at a rate of 35°/s. The dot expansion was rendered over a sequence of video frames by incrementally increasing its diameter and redrawing it at the center of the screen on each frame. Sweeping dark dot (Condition 2): In this condition, the stimulus combined motion with a decrease in intensity: A black dot first appeared at one of the four screen corners and moved in a straight trajectory toward the screen center at a constant angular speed of 10°/s. After reaching the center, the dot remained stationary and underwent the same radial expansion (4° to 40°) as in Condition 1. The light intensity of the stimuli did not exceed 100 lux as measured with a lux-meter (RS PRO ILM 1332A). Stimulus onset and timing were synchronized using a photodiode placed at a fixed position on the screen and modulated concurrently with the main stimulus.

#### Tetrode and Neuropixel recordings with freely moving mole-rats

Tetrode recordings were performed using 8-tetrodes made from platinum-iridium wires enclosed in microdrives (Neuralynx, VersaDrive-8). Tetrodes were gold plated using NanoZ with resultant impedance values < 200 kΩ. Neural activity was recorded using the Neuralynx Data Acquisition System (Digital Lynx SX). The LFP data was acquired continuously on all four channels of the tetrodes and filtered with a finite impulse response (FIR) filter in the range of 1 to 500 Hz. Spike data was recorded on all four channels for the tetrodes in continuous format and spike triggered format using a threshold value of 50 µV and bandpass filtered using a FIR filter in the range of 600 to 5,000 Hz. Data was recorded from two animals and 1 to 4 sessions on multiple days. During the recordings, the animals were exploring a circular track (track width: 5 cm, outer diameter of the circle: 80 cm). The tetrodes were lowered by 30 µm between consecutive recording days.

In freely moving experiments with the Neuropixels 2.0 probe, the animals were placed in a circular track (Figure 3A, details above) that they were allowed to explore, and the neural activity was collected at 30 kHz using the Neuropixels data-acquisition system as described for the head-fixed recordings above.

#### Video recording and animal tracking system for freely moving animals

Videos of the animals in the arena were recorded using a Basler ace U acA1300-75gm camera in IR light (peak wavelength 880 nm, bandwidth 50 nm, Kingbright KP-2012SF4C) at 30 fps. Infrared LEDs with two different peak wavelengths (880 nm and 940 nm, Kingbright KP-2012SF4C and KP-2012F3C) were mounted on the implanted Versa drive/Apollo drive to enable tracking with DeepLabCut (Version 2.3.9) (Mathis et al., 2018).

#### Neuropixels data analysis

We used SpikeInterface (Buccino et al., 2020) and Kilosort 4 (Pachitariu et al., 2024) to analyze Neuropixels data. Neuropixels 1.0 LFP data were loaded into SpikeInterface and filtered for different frequency bands (Theta: 4-8 Hz, Alpha: 8-15 Hz, Beta: 15-30 Hz, and Gamma: 30-100 Hz). LFP power in each frequency band was calculated by Welch’s method over 1-second intervals and averaged across ten intervals. Current source density analysis was performed using broad-band LFP (0.5-250 Hz) and its spatial second derivative. The power of current source density was computed using Welch’s method, similar to other LFP powers.

To analyze the spike data, we preprocessed the raw data by applying a high-pass filter at 400 Hz and common reference averaging in SpikeInterface. We then used Kilosort 4 with default parameters for spike sorting. We computed the quality metrics using SpikeInterface and filtered the detected clusters by applying thresholds on ISI violations (0.4) and amplitude cutoff (0.2). This automated clustering was followed by manual curation to further split or merge clusters and remove noise clusters.

#### Tetrode data analysis

The tetrode data was analyzed using custom MATLAB codes. The LFP data was down-sampled from 32 kHz to 250 Hz. The instantaneous speed was calculated using the change in distance between individual frames that were 33.33 ms apart. The spectrogram was plotted by calculating a short-time Fourier transform of the signal with a moving window of 1,000 sample points with a 50% overlap between windows. The power spectral density was computed by the Welch method using a Hamming window of 8 segments with 50% overlap. The data was converted to dB by normalizing the power to the maximum power during the session. The range of theta frequency was determined using the average value of the upper and lower bounds determined by applying the full width at half maximum on the kernel density estimate curve of the power spectrum. The LFP data were filtered using a Butterworth filter (4^th^ order) in the range of 3 to 6 Hz. The instantaneous phase of theta was determined using Hilbert transform.

#### Statistical analysis

Statistics were performed in GraphPad Prism (Version 10.3.1). Data were tested for normality using Q-Q plots, followed by appropriate parametric or nonparametric statistical tests.

### Histology

#### Perfusion

At the end of the electrophysiological recordings, the animals were euthanized with a high dose of ketamine (300 mg/kg) and xylazine (30 mg/kg). The animals were perfused with 100 ml of PBS supplemented with 10 U/ml of heparin, followed by 100 ml of 4% paraformaldehyde in 0.1 M PB (PFA) solution at a flow rate of 7 ml/min. The head was postfixed in 4% PFA for 24 h, after which the brains were dissected and stored in PBS supplemented with 0.02% sodium azide at 4°C.

#### Tissue clearing and light sheet microscopy

To achieve optical transparency for three-dimensional imaging, we used a modified iDISCO^+^ tissue-clearing protocol (Renier et al., 2016) (detailed protocol in Table S2). In brief, the brains were dehydrated in increasing concentrations of methanol (10% to 100%) in phosphate-buffered saline (PBS). After dehydration, tissues were delipidated in a 66%/33% dichloromethane (DCM)/methanol solution followed by 100% DCM. Finally, the samples were incubated in dibenzyl ether (DBE) to achieve refractive-index matching. All steps were performed in opaque 50 ml centrifuge tubes (Greiner Bio-One) or 5 ml reaction tubes (Eppendorf) made from polypropylene (PP) at room temperature (RT) on a rotator to ensure uniform exposure to clearing solutions. Cleared tissues were stored in DBE at RT until imaging. Imaging was performed in DBE using a commercial light-sheet microscope (Miltenyi Biotec Ultramicroscope 2). Autofluorescence was recorded at 488 nm excitation wavelength, while the fluorescence signal from the CM-DiI dye labelled electrode track was visualized with a 561 nm excitation wavelength. Imaging was performed using infinity-corrected optics and a 1.1x MI PLAN objective (Miltenyi Biotec), resulting in a pixel size of 6.57 µm in the x-y plane and 8 µm along the z-axis. The dual-channel scanning allowed us to map the brain’s structural organization and the electrode tracks with high resolution and contrast. Image processing was performed in Arivis Vision 4D (Zeiss) and Fiji ImageJ 1.53t (Schindelin et al., 2012).

#### Reversed tissue clearing (reDISCO)

We developed a protocol to reverse the iDISCO+ clearing which we refer to as reDISCO (Table S3). Due to the light sensitivity of the CM-Dil tracer, light exposure should be minimized. Cleared brains stored in DBE were rehydrated in two steps of 100% DCM, followed by two washes in 100% MeOH before being rehydrated in decreasing concentrations of methanol in PBS (80% to 10%). All steps were performed in opaque 50 ml centrifuge tubes (Greiner Bio-One) or 5 ml reaction tubes (Eppendorf) made from PP at RT on a rotator. The rehydrated brains were stored in PBS + 0.02% sodium azide at +4°C until sectioning.

To prepare for vibratome (Leica VS1200S) sectioning, all liquid at the brain surface was removed carefully with a soft tissue (Kimwipe) and brains were embedded in 4% low melt agarose in PBS. After hardening, the agarose block was fixed on the vibratome stage with super glue. 100 µm sections were obtained at 1.00 mm/s speed and 1.00 mm amplitude, collected in PBS, mounted onto SuperFrost Ultra Plus slides, dried for 90 minutes at RT and stained.

Sections were washed in 1x PBS and stained with fluorescent Nissl (Neurotrace 640/660) 1:1000 in 1x PBS / 0.3% Triton-X 100 in a humidified chamber for one hour. After staining, slides were washed in 1x PBS, nuclei were counterstained with DAPI in PBS (1:2000), washed with PBS and coverslipped with ROTIMount FluorCare. All staining steps were performed at RT, and the stained slides were stored at +4°C until imaging at an automated epifluorescent slide scanner (Evident SLIDEVIEW VS200) with a 20x objective at an excitation wavelength of 550 nm for the Neuropixel tracks (CM-Dil), 630 nm for Nissl Neurotrace and 405 nm for DAPI.

#### Tetrode track reconstruction

To reconstruct the tetrode tracks after transcardial perfusion, PFA-fixed brains were cryoprotected in 30% sucrose in PBS, and 40-micron-thick sections were cut using a frozen-stage rotary microtome (Microm HM 340). The sections were mounted on SuperFrost Ultra Plus slides and rehydrated through graded ethanol concentrations and PBS washes. The sections were then stained using 1% Cresyl violet solution, followed by differentiation and dehydration using ascending ethanol concentrations and xylene. Finally, the slides were coverslipped with DPX new (Sigma-Aldrich) for imaging. Slides were scanned with a 20x objective using an automated slide scanner (Evident SLIDEVIEW VS200).

#### Micro-computed-tomography (µCT) and rendering

To assess the structure of the skull at the area which is subject to surgery (i.e., the Os parietale and Os frontale) and the muscles covering this area (i.e., the temporalis muscle), µCt data were obtained from several mole-rat specimens. For *F. anselli,* data was acquired from five specimens (IDs: Fa_30972, Fa_D9C67, Fa_FE839, Fa_FDBFFA, FA93). For comparison, µCT data was also acquired for *Fukomys micklemi* (ID: 7527), *Heterocephalus glaber* (ID: NMR3), and *Mus musculus* (ID: 314). Specimens were fixed by transcardial perfusion (see above) or drop-fixation in 4% PFA. For contrast enhancement of soft tissues, animals or heads were immersed in 1% Lugol solution (N052.2; Carl-Roth) for 4 weeks up to 1 year, with 2-3 changes of the solution. For µCT scanning, the samples were mounted in PBS in an X-radia Versa 620 (Zeiss). Details on the scanning parameters are given in Table S4. Image stacks were rotated and cropped, and the thickness of the Os frontale and Os parietale as well as of the sagittal ridge, were measured (see Figure S1) in Fiji ImageJ 1.53t (Schindelin et al., 2012). The skull and the temporalis muscles were segmented in 3D Slicer 5.6.1 (Kikins et al. 2014). Rendering was performed using Meshlab v2022.02 (Cignoni et al., 2008) and Blender 2.77 (https://www.blender.org/).

### Stereotaxic Brain Atlas

#### Tissue preparation

We developed a stereotaxic brain atlas for *Fukomys anselli* mole-rats to facilitate accurate targeting of brain regions during electrophysiological recordings. The atlas was created using data from two adult male mole-rats (83 weeks old, 66.3 g; 94 weeks old, 109.6 g). The animals were placed in a stereotaxic frame under surgical anesthesia, and track marks were placed by drilling through the skull and driving a microneedle into the brain at eight bilateral positions spaced 2 mm apart using Bregma as a reference point (Figure S5A). The distance between these marks was divided by the number of sections between them to calculate the corrected section thickness. The track marks level with the anterior-posterior axis relative to Bregma were designated as the 0 mm reference points. After the track marks were placed, the animals were euthanized with a high dose of ketamine (300 mg/kg) and xylazine (30 mg/kg) and perfused with heparinized PBS followed by 4% PFA in 0.1 M PB. The brains were extracted and post-fixed overnight in 4% PFA at 4°C. For cryoprotection, the brains were incubated in 30% sucrose in PBS until they sank. The brains were then sectioned coronally at 40 µm using a frozen-stage microtome (Microm HM 340), using the track marks to ensure proper alignment along the anterior-posterior axis. Every second section was mounted and stained using 1% Cresyl violet solution, followed by differentiation and dehydration using ascending ethanol concentrations and xylene. Finally, the slides were mounted with DPX new (Sigma-Aldrich) for imaging. Slides were scanned with a 20x objective using an automated slide scanner (Evident SLIDEVIEW VS200).

#### Atlas Construction and Alignment

Although the brain sections were cryosectioned at 40 µm, the actual thickness did not correspond to this value in live tissue due to tissue shrinkage (∼25%) caused by perfusion and sucrose incubation. Therefore, all measurements were corrected to account for this reduction. Moreover, any sections lost during slicing or mounting were recorded, and their distances were interpolated from the track marks created during slicing. This method ensured accurate alignment and consistency of stereotaxic coordinates across the atlas. The atlas was constructed by aligning sections from one mole-rat, using Bregma and Lambda as stereotaxic reference points. Sections from the second mole-rat were incorporated to fill gaps and ensure continuity. Lost sections were accounted for by interpolating distances based on track marks and ensuring consistent spacing across slices. The atlas was annotated by comparing the Nissl-stained sections with existing brain atlases, including one for *Fukomys anselli* (Dollas et al., 2019), Paxinos and Franklin’s mouse brain atlas (Paxinos & Franklin, 2019), Paxinos and Watson’s rat brain atlas (Paxinos & Watson, 2006), and the Allen Mouse Brain Atlas (Wang et al., 2020). We included only regions with clearly delineated anatomical features. We used Affinity Designer (Serif Europe Ltd) to create the atlas plates. Each plate features a coronal section on the left, annotated with anatomical abbreviations listed on the right. A sagittal view of a schematic mole-rat brain is displayed at the top right, indicating the position of the coronal section relative to Bregma (Figure S5B).

#### CO_2_ measurements

Carbon dioxide concentrations were measured using a CO**_2_** sensor (Vernier CO2-BTA) placed in a gas sampling bottle with cage air collected by suction. The Logger Lite app (Vernier) was used to record the data. The CO**_2_** concentration at different locations in the room and cages was measured by collecting the air in the bottle. We collected multiple samples of air from the i) animal housing room, ii) home cage, iii) 100 cm long and 5 cm wide cylindrical tunnels in the home cage, iv) nest in the home cage, and v) the enclosed recovery box placed inside the home cage (Figure S4B).

#### Sequencing of KCC2 fragments

We extracted mRNA from snap-frozen (liquid nitrogen) brain tissue of two *Fukomys anselli* specimens (ZYMO Research, Quick-RNA Miniprep plus kit) and prepared cDNA libraries (Qiagen Quantitect reverse transcription kit) according to the manufacturer’s instructions. Based on the *Fukomys anselli* KCC2 sequence taken from our transcriptomic dataset of various tissues, Primers G0493 (TGGCCCAGATGGACGACAACAGC) and G0494 (TTCAAGTTCTCCCACTCCGGCTTC) were then used to amplify a fragment of the KCC2 transcript via PCR with KOD Hot Start Polymerase (Sigma-Aldrich) using the pooled brain cDNA libraries as template. Forty cycles were performed according to the following protocol: 30 s at 94 °C, 20 s at 56 °C, and 40 s at 68 °C. The PCR products were purified from the agarose gel (Nucleospin Gel & PCR Clean-up, Macherey-Nagel) and Sanger sequenced (Eurofins) using the sequencing primer G0497 (AGCATCCAGATGAAGAAGGACCTG).

