## Supplemental information for "Neuronal recordings in head-fixed and freely-moving mole-rats"

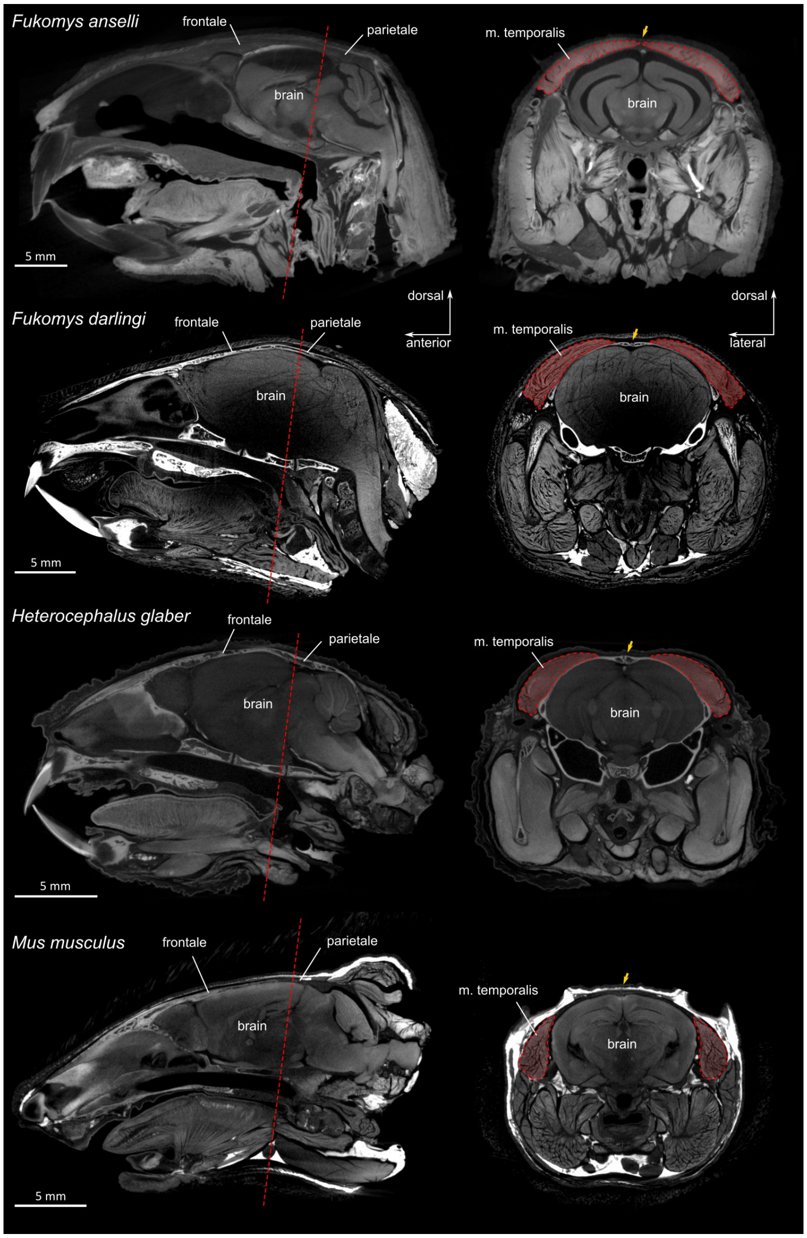


**Figure S1. Micro-CT scans reveal the increased dorsal muscle coverage of the frontal and parietal skull bones (i.e., the area of the craniotomy) in three social African mole-rats compared to the mouse.** Comparison of the skull (white) and temporal muscles (red) covering the skull's Os frontale and Os parietale in different Bathyergidae (from top to bottom: *Fukomys anselli, Fukomys darlingi, Heterocephalus glaber*) and laboratory mouse (bottom: *Mus musculus, C57BL/6J*). Left: sagittal section through the head midline with dotted lines indicating the cutting plane of cross sections. Right: cross-sections as indicated on the left with the temporalis muscle marked in red and the midline suture marked by a yellow arrow.

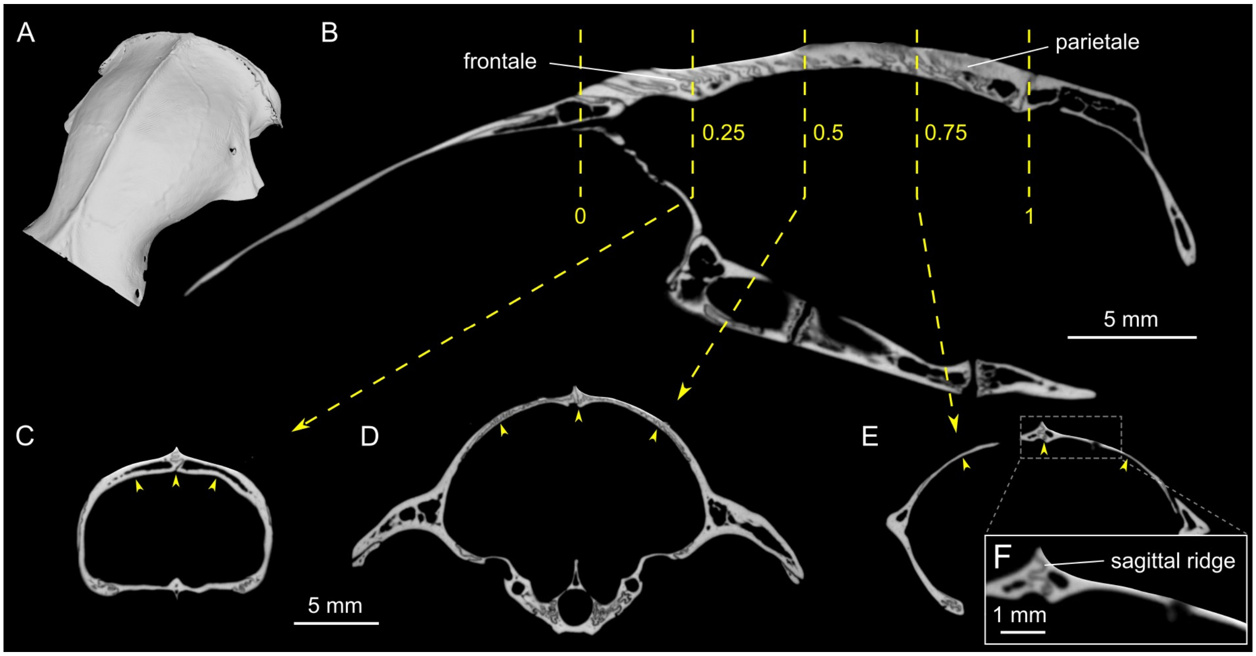


**Figure S2. Measurement of skull structure and thickness in *Fukomys anselli* based on µCT data. A.** Surface rendering of the dorsal cranium with the prominent midsagittal ridge. **B.** Sagittal section of the head through the midline, indicating places of cross-sections shown in C-E. **C.** Cross-section at 0.25 length of Os frontale and Os parietale. **D.** Cross-section at 0.5 length of Os frontale and Os parietale. **E.** Cross-section at 0.75 length of Os frontale and Os parietale, revealing an area where the skull is very thin. **F.** Detail of Os parietale with sagittal ridge in E.

**Table S1.** Skull bone thickness in *F. anselli* (n = 4) based on µCT data. All measurements in µm.

| **Specimen** | **0.25 L** | **0.25 R** | **0.25 M** | **0.5 L** | **0.5 R** | **0.5 M** | **0.75 L** | **0.75 R** | **0.75 M** |
|---|---|---|---|---|---|---|---|---|---|
| Fa_DBFFA | 570 | 546 | 1,112 | 416 | 389 | 1,083 | 126 | 157 | 1,116 |
| Fa_30972 | 330 | 378 | 1,532 | 440 | 466 | 860 | 158 | 192 | 1,780 |
| Fa_D9C67 | 250 | 275 | 1,277 | 362 | 352 | 1,037 | 130 | 181 | 1,313 |
| Fa_FE839 | 344 | 401 | 980 | 465 | 544 | 537 | 185 | 200 | 664 |
| **Mean** | **373** | **400** | **1,225** | **420** | **437** | **879** | **150** | **183** | **1,218** |
| **SD** | **119** | **97** | **206** | **38** | **74** | **214** | **24** | **16** | **401** |

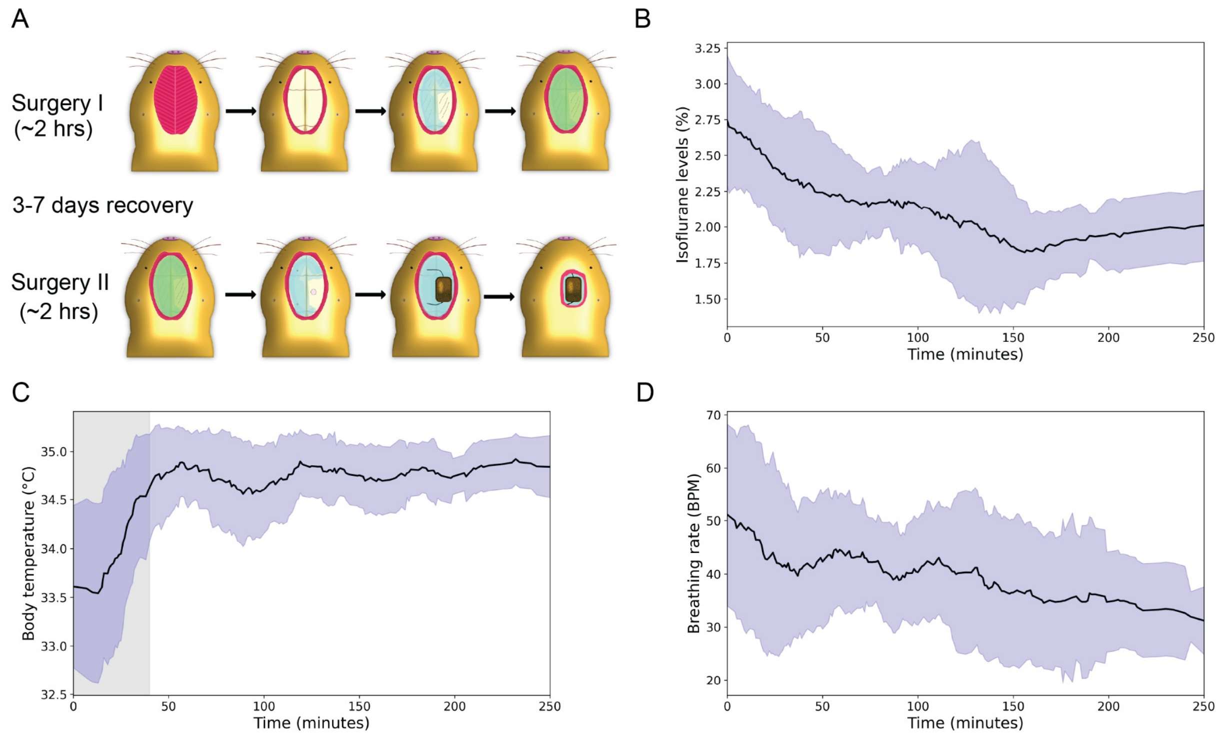


**Figure S3: Two-part surgery and vital parameters of *Fukomys* mole-rats under isoflurane anesthesia. A.** Two-step surgical procedure. In the first surgery, the muscles covering the skull are carefully removed, and the skull is cleaned, coated with a base layer of dental composite, and covered with Kwik-cast. After 3-7 days of recovery, Kwik-cast is removed, the craniotomy is performed, the implant is secured with dental composite, and the wound is closed with tissue glue and/or surgical sutures. **B.** Isoflurane levels during successful surgeries. After induction at 3%, the concentration was maintained at around 2% throughout the surgery (n = 18 surgeries; Mean ± SD). **C.** Body temperature during successful surgeries. The closed-loop system was set to 34.9°C, but the body temperature of *Fukomys* mole-rats showed notable fluctuations (n = 18 surgeries, Mean ± SD). The shaded area indicates the warm-up phase of the probe after rectal insertion. **D.** Breathing rate during successful surgeries. The breathing rate was highly variable between specimens and decreased over time (n = 18 surgeries, Mean ± SD).

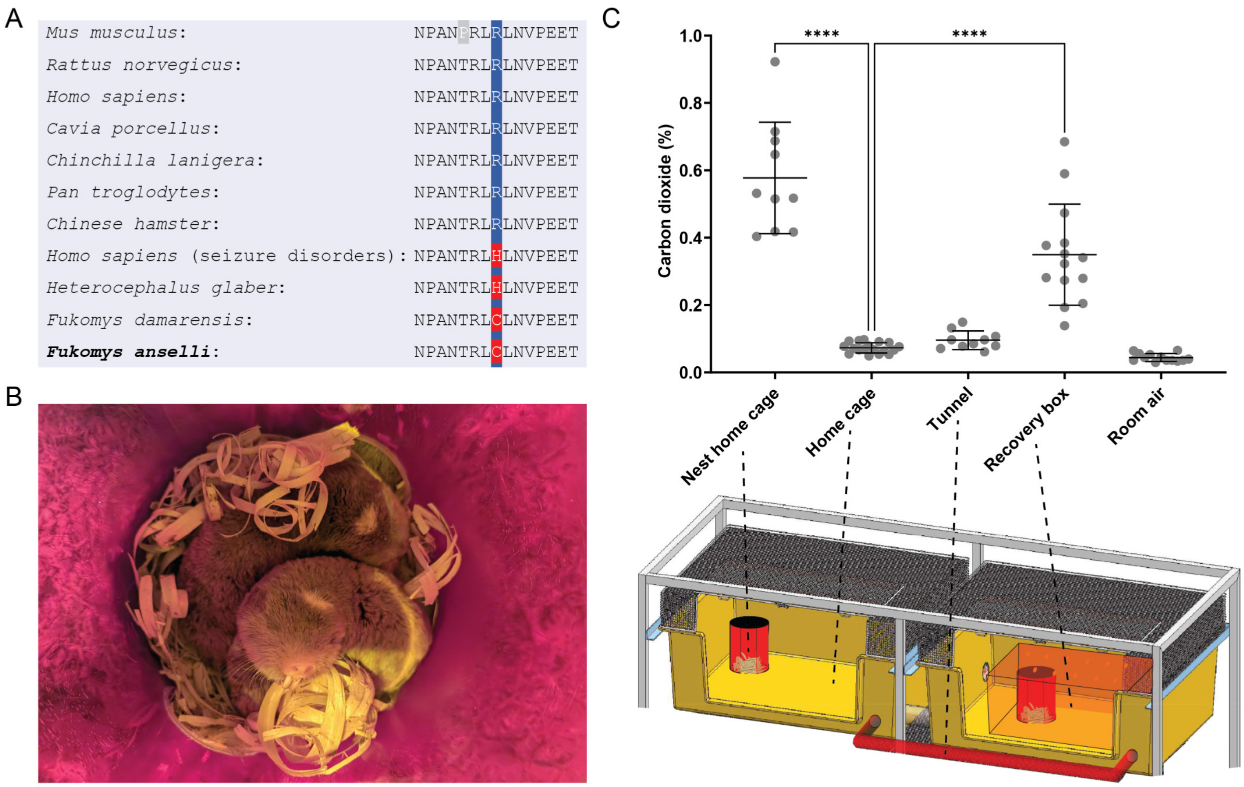


**Figure S4. A seizure-related KCC2 variant and elevated $CO_2$ concentrations in the nest boxes and post-surgery recovery environment. A.** Alignment of the amino acid sequences encoded within exon 22 of KCC2 orthologs in rodent and primate species. Sanger sequencing of PCR products from brain cDNA revealed that the highly conserved amino acid arginine (R) at position 952 (in human KCC2b; NCBI: NP_065759.1) carries a cysteine (C) in the Ansell's mole-rat *Fukomys anselli*. The exact position also carries a cysteine in the Damaraland mole-rat (*Fukomys damarensis*; NCBI: XP_010624588.1), and a histidine (H) in the naked mole-rat (*Heterocephalus glaber*; NCBI: XP_004858176.1). The latter mutation of the gene encoding KCC2 (R952H) was found in an Australian family with febrile seizures (Puskarjov et al., 2014) and in a cohort of Canadian patients with idiopathic generalized epilepsy (Kahle et al., 2014). R952H is also related to human autism spectrum disorders and schizophrenia (Merner et al., 2015). All sequences except for *Fukomys anselli* were taken from the in-silico analysis presented in Zions et al. (2020). **B.** Mole-rat family huddling in a nest box. Note that parts of the bedding were removed to take the photo. **C.** Carbon dioxide concentrations in different parts of the mole-rat home cages. The animals are housed in cages connected with tunnels and provided with nest boxes in which $CO_2$ builds up to concentrations approximately 10 times higher than in the room atmosphere. Providing the animals with similar $CO_2$ concentrations in a minimally ventilated recovery box significantly reduced the prevalence of seizures during the recovery period. One-Way ANOVA with Tukey's post hoc multiple comparisons. ****: $p < 0.0001$

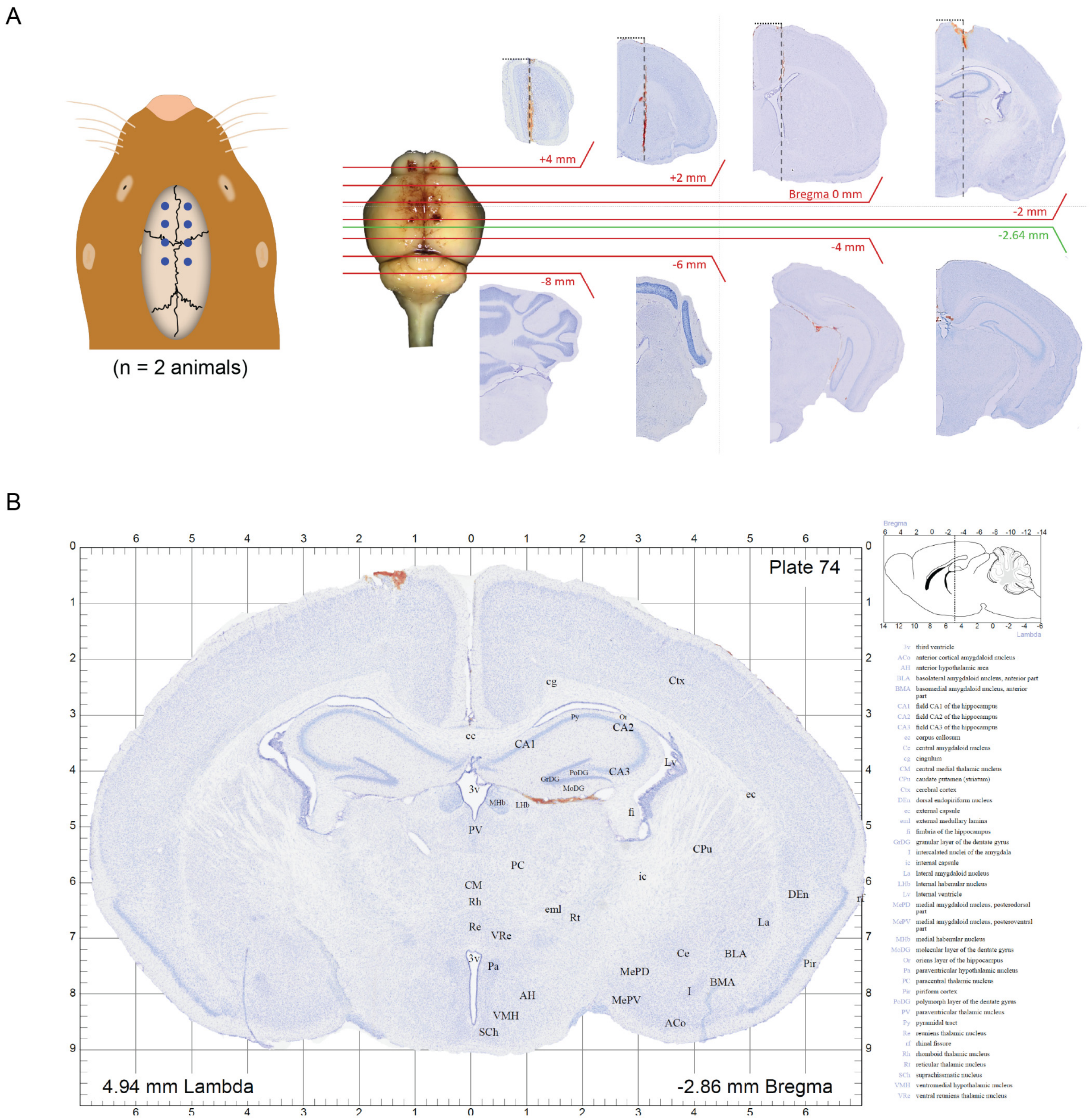


**Figure S5. A stereotaxic brain atlas of *Fukomys anselli.* A.** To obtain atlas coordinates, two anesthetized mole-rats were placed in a stereotaxic frame, and track marks were placed by drilling through the skull and driving a microneedle into the brain at eight bilateral positions spaced 2 mm apart using Bregma as a reference point. The distance between these marks was divided by the number of sections between them to calculate the corrected section thickness. The track marks level with the anterior-posterior axis relative to Bregma were designated as the 0 mm reference points. **B.** Example plate of the stereotaxic atlas. Each plate features a coronal section on the left, annotated with anatomical abbreviations. A sagittal view of the mole-rat brain is displayed at the top right, indicating the position of the coronal section relative to Bregma. Complete abbreviations are listed on the right of the plate. The full atlas with 175 plates is available at doi.org/10.17617/3.UNDKRO.

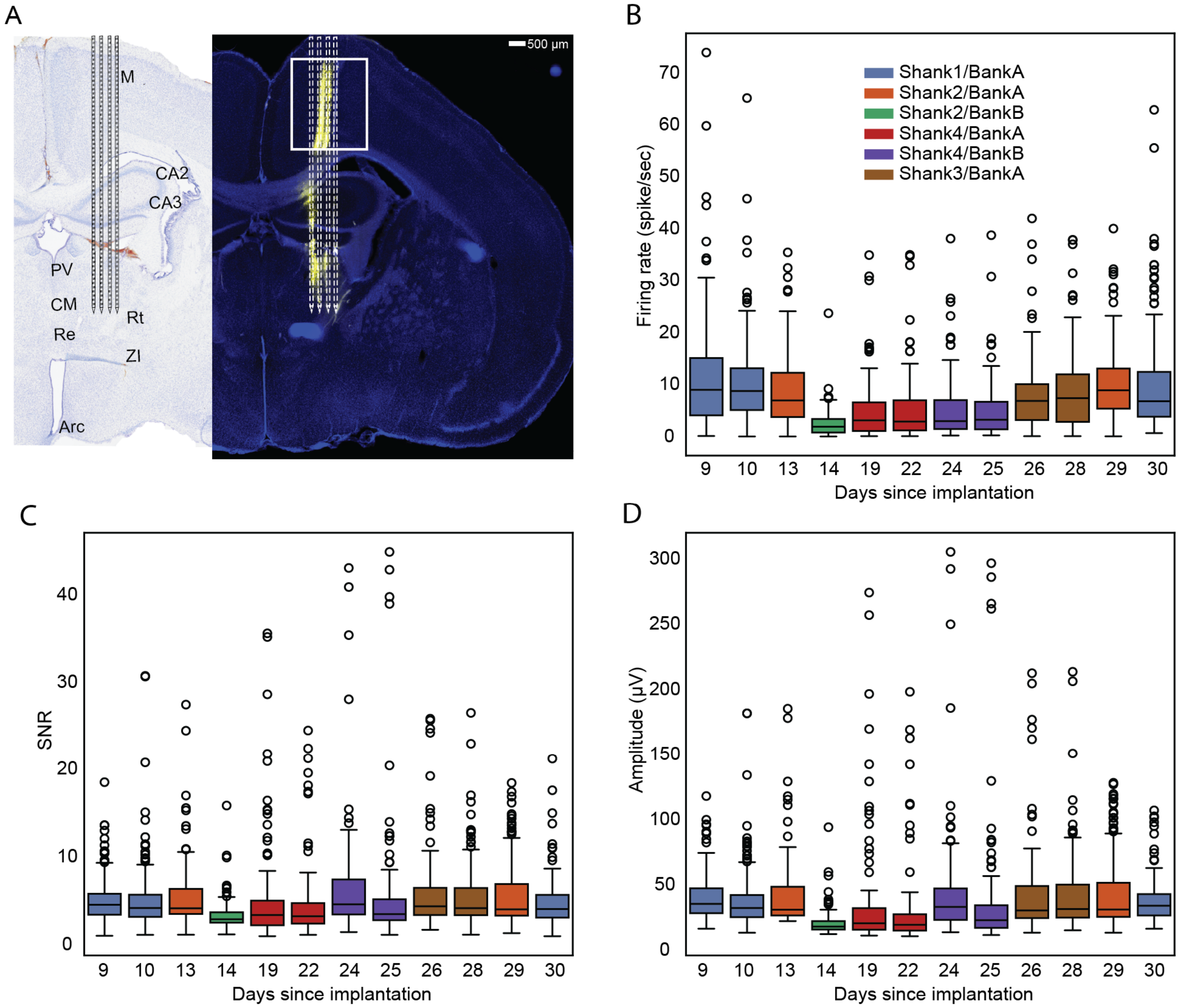


**Figure S6. Quality metrics of recorded units using a Neuropixels 2.0 probe in a freely moving Ansell's mole-rat. A.** Histological reconstruction of the NPX 2.0 probe track with CM-DiI (yellow) in 100 µm vibratome sections stained with DAPI after tissue clearing and a reverse iDISCO protocol next to the corresponding plate from the stereotaxic Ansell's mole-rat brain atlas. **B.** Firing rate of recorded units over time. Each experiment included recorded data from 384 channels in a single shank. The color indicates the shank and bank used for data collection. **C. and D.** The box plots show the signal-to-noise ratio (SNR) and amplitude (µV) of the recorded units over time, respectively. The color code is the same as in B.

**Table S2.** Brain clearing protocol (modified iDISCO$^+$ protocol (Renier et al., 2016)). DBE: Dibenzyl ether; DCM: Dichloromethane; PBS: Phosphate-buffered saline (0.1 M); RT: room temperature. The individual incubation times may be extended. Samples should be imaged within two weeks after clearing to ensure a high signal-to-noise ratio.

| Day | Incubation Time | Solution | Conditions |
|---|---|---|---|
| 1 | 12 h | 10% Methanol in PBS | RT, on rotator |
| 1 | 12 h | 20% Methanol in PBS | RT, on rotator |
| 2 | 12 h | 30% Methanol in PBS | RT, on rotator |
| 2 | 12 h | 40% Methanol in PBS | RT, on rotator |
| 3 | 12 h | 50% Methanol in PBS | RT, on rotator |
| 3 | 12 h | 60% Methanol in PBS | RT, on rotator |
| 4 | 12 h | 70% Methanol in PBS | RT, on rotator |
| 4 | 12 h | 80% Methanol in PBS | RT, on rotator |
| 5 | 12 h | 100% Methanol | RT, on rotator |
| 5 | 18 h | 100% Methanol | RT, on rotator |
| 6 | 1 h | 66/33% DCM/Methanol | RT, on rotator |
| 6 | 24 h | 66/33% DCM/Methanol | RT, on rotator |
| 7 | 24 h | 66/33% DCM/Methanol | RT, on rotator |
| 8 | 1 h | 100% DCM | RT, on rotator |
| 8 | 24 h | 100% DCM | RT, on rotator |
| 9 | 24 h | 100% DCM | RT, on rotator |
| 10 | 24 h | DBE | RT (no rotation) |
| 10 | Store until imaged | DBE | RT (no rotation) |

**Table S3.** Reverse brain-clearing protocol (reDISCO). This protocol describes how to rehydrate DISCO-cleared brains for vibratome sectioning. DBE: Dibenzyl ether; DCM: Dichloromethane; PBS: Phosphate-buffered saline (0.1 M); RT: room temperature. The individual incubation times may be extended.

| Day | Time | Solution | Conditions |
|---|---|---|---|
| 1 | 1 h | 100% DCM | RT, on rotator |
| 1 | 23 h | 100% DCM | RT, on rotator |
| 2 | 24 h | 66% DCM/33%Methanol (stock) | RT, on rotator |
| 3 | 24 h | 66% DCM/33%Methanol (stock) | RT, on rotator |
| 4 | 24 h | 100% Methanol | RT, on rotator |
| 5 | 12 h | 100% Methanol | RT, on rotator |
| 5 | 12 h | 80% Methanol/PBS | RT, on rotator |
| 6 | 12 h | 70% Methanol/PBS | RT, on rotator |
| 6 | 12 h | 60% Methanol/PBS | RT, on rotator |
| 7 | 12 h | 50% Methanol/PBS | RT, on rotator |
| 7 | 12 h | 40% Methanol/PBS | RT, on rotator |
| 8 | 6 h | 30% Methanol/PBS | RT, on rotator |
| 8 | 6 h | 20% Methanol/PBS | RT, on rotator |
| 8 | 18 h | 10% Methanol/PBS | RT, on rotator |
| 9 | 24 h | PBS + 0.02% sodium azide | RT, on rotator |
| 10 | Embedding in 4% agarose and vibratome sectioning | | |

**Table S4.** Specimens (IDs) and parameters used for µCT scanning.

| Species | Specimen (ID) | Lugol | Fig. | Voxel size (µm) | Voltage (kV) | Power (W) | Exposure (s) | Projections |
|---|---|---|---|---|---|---|---|---|
| *M. musculus* | 314 | 29.11.2023 - 26.11.2024, changed 3x = 1 year | CT | 17.66 | 130 | 14 | 0.4 | 4,501 |
| *F. anselli* | FA24_1593 | 1% Lugol for 216h 10% Lugol for 264h | CT | 17.17 | 140 | 15 | 0.1 | 4,501 |
| *F. anselli* | FAFE839 | none | Tab. S1 | 28.28 | 120 | 12 | 0.5 | 4,501 |
| *F. anselli* | FAD9C67 | none | Tab. S1 | 26.44 | 120 | 13 | 1.0 | 4,501 |
| *F. anselli* | FA30972 | none | Tab. S1 | 20.00 | 120 | 12 | 0.3 | 4,501 |
| *F. anselli* | FDBFFA | none | Tab. S1 | 23.81 | 120 | 12 | 0.2 | 4,501 |
| *F. darlingi* | FD2, 5361m | 07.11.2023-22.11.2023, changed 3x = 1 year | CT | 17.16 | 120 | 13 | 0.5 | 3,001 |
| *H. glaber* | NMR2 | 27.10.2023-08-01.2024, changed 1x = 10 weeks | CT | 15.10 | 130 | 14 | 0.5 | 4,501 |
| *H. glaber* | NMR3 | 27.10.-27.11.2023, changed 1x = 4 weeks | 3D | 13.40 | 130 | 14 | 0.5 | 4,501 |

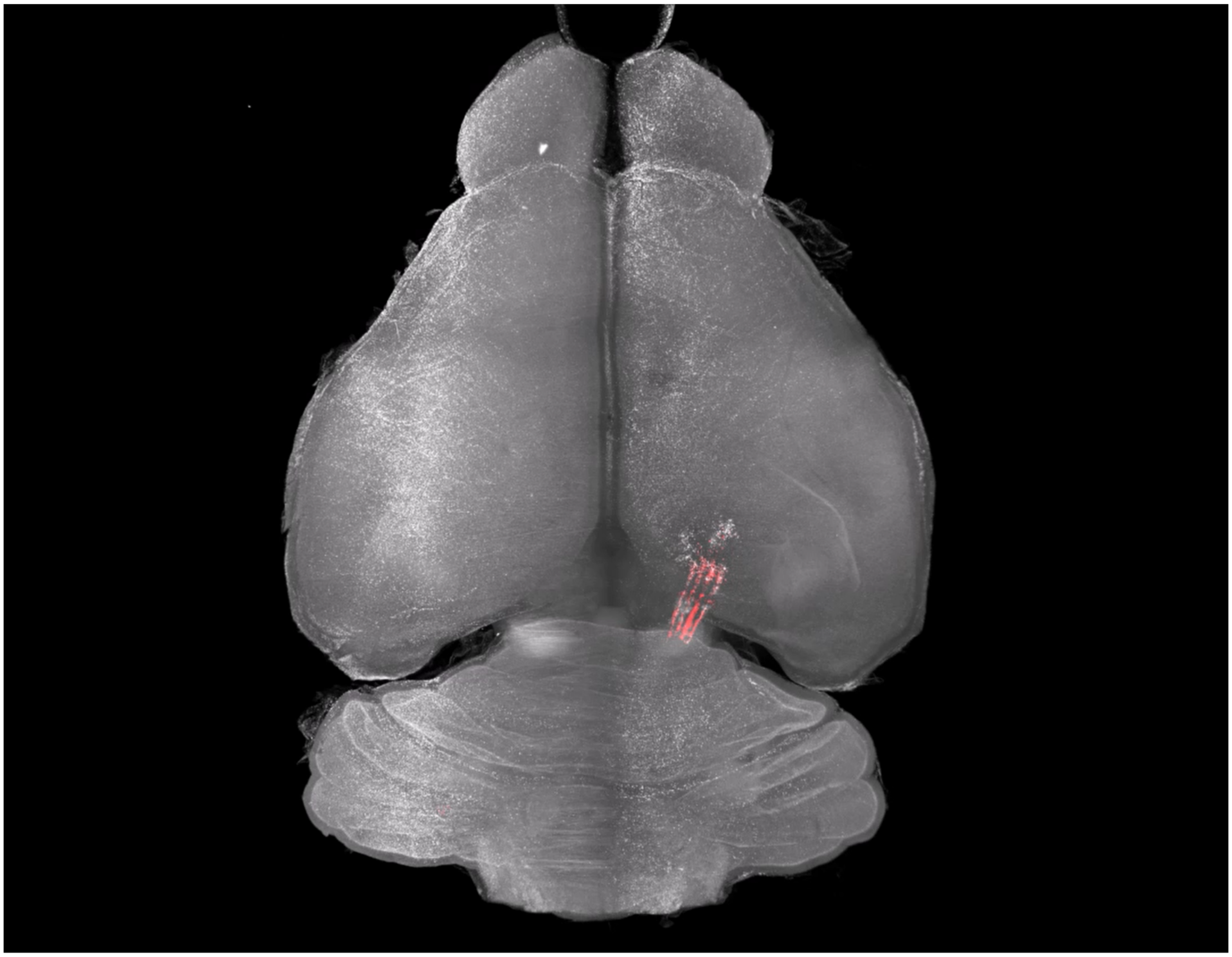

**Video S1.** 3D-reconstruction of the tracks of a NPX2.0 probe in the Ansell's mole-rat brain based on tissue clearing and light sheet imaging. The video was created in Arivis vision 4D and can be viewed here.
